# Parkinson’s disease linked LRRK2 G2019S drives oxidative nuclear DNA damage and PARP1 hyperactive signaling

**DOI:** 10.64898/2026.02.27.708379

**Authors:** Jennifer Liu, Claudia P. Gonzalez-Hunt, Tara Richbourg, Ivana Barraza, Carolyn Chen, Carlos Montes, Lingyan Ma, Ruihua Cao, Vishruth Hanumaihgari, Natalie R. Gassman, Elise Fouquerel, Laurie H. Sanders

## Abstract

*LRRK2* mutations are the most common cause of autosomal-dominant Parkinson’s disease (PD), with G2019S linked to both familial and sporadic PD. Although LRRK2-mediated mitochondrial DNA damage is implicated in PD, the contribution of nuclear DNA damage is less understood. Using CRISPR/Cas9-generated LRRK2^G2019S/G2019S^ knock-in cells, we discovered increased sensitivity to oxidative and alkylating DNA-damaging agents compared to wild-type, consistent with compromised tolerance/repair of lesions processed by base excision repair (BER). The oxRADD assay revealed elevated endogenous oxidative nuclear base damage in *LRRK2* mutant cells. Concomitantly, PARP1-dependent poly(ADP-ribose) (PAR) levels were markedly increased, with chromatin enrichment of PARP1 and BER factors (XRCC1, DNA ligase III) only in LRRK2^G2019S/G2019S^ cells, indicating BER initiation, without successful resolution. LRRK2^G2019S/G2019S^ cells displayed synthetic lethality with PARP-trapping inhibitors (olaparib) but tolerated PARP1 knockdown, suggesting cytotoxicity from stabilized PARP-DNA complexes rather than loss of catalytic activity. The SOD/catalase mimetic EUK-134 abrogated LRRK2 G2019S-dependent PAR accumulation, whereas the mitochondrial complex I inhibitor rotenone exacerbated PAR levels, linking reactive oxygen species (ROS) to BER dysfunction and PARP1 hyperactivation. Overall, we have identified a ROS-dependent PARP1 hyperactivation pathway that underlies LRRK2 G2019S-associated cellular vulnerability.

## INTRODUCTION

Parkinson’s disease (PD) is an age-associated neurodegenerative movement disorder affecting 1–2% of people over 65, with prevalence rising as populations age (1–4). Clinically, PD is characterized by bradykinesia, rigidity, tremor, and gait disturbance, and non-motor symptoms are also common (5). Pathology principally involves degeneration of dopaminergic neurons in the substantia nigra pars compacta and accumulation of misfolded and aggregated proteins such as alpha-synuclein (6,7). Although most PD cases are sporadic, ∼10–20% are familial (8–10). Leucine-rich repeat kinase 2 (LRRK2) is the most frequent cause of autosomal dominant late-onset PD; pathogenic *LRRK2* variants, notably Gly2019Ser (G2019S), accounts for ∼4% of familial and ∼1% of sporadic PD cases, with higher prevalence in some populations (11,12). Because LRRK2-linked PD clinically closely resembles sporadic disease and LRRK2 is implicated in sporadic PD even in the absence of a mutation, studying LRRK2 can provide insights into broader PD mechanisms (13–15). LRRK2 encodes a large multidomain protein with kinase and GTPase activities; PD pathogenic variants increase kinase activity, motivating therapeutic efforts to target LRRK2 (11,16,17). LRRK2 phosphorylates Rab GTPases and, together with its substrates, has been implicated in a variety of cellular functions, including but not limited to autophagy/lysosomal and mitochondrial function (11).

Mitochondrial dysfunction is a central mediator of PD pathogenesis, and mounting evidence ties LRRK2 to mitochondrial homeostasis (18). Pathogenic *LRRK2* variants, particularly G2019S, have been associated with impaired mitophagy and dynamics, altered bioenergetics, and increased mitochondrial DNA (mtDNA) damage (18). mtDNA damage or lesions (e.g., 8-oxo-dG, abasic sites) and mutations (deletions, point changes, transversions) have been reported in experimental PD models and postmortem brains derived from patients with PD (19–25). We and others have observed LRRK2 G2019S–dependent mtDNA damage in iPSC-derived neurons, lymphoblastoid cells, LRRK2 G2019S knock-in mice, G2019S LRRK2 PD patient derived peripheral blood mononuclear cells, and LRRK2 G2019S knock-in HEK293 cells (26–30). Although the mechanisms linking LRRK2 to mtDNA integrity are incompletely defined, mitochondrial defects increase reactive oxygen species (ROS), which compromise bioenergetics and further cause oxidative damage to lipids, proteins, and nucleic acids; indeed, multiple biochemical, genetic, histopathologic, and biomarker studies implicate oxidative stress in PD (21,24,31–34).

Age is the strongest risk factor for PD; aging is associated with increased oxidative stress and reduced DNA repair capacity, processes linked to neurodegenerative diseases including PD (35–38). Unrepaired DNA lesions can disrupt fundamental cellular processes, and accumulation or persistence of DNA damage can promote genomic instability, cellular senescence, or cell death (39,40). In both the mitochondria and nucleus, the major pathway for repair of oxidative base lesions, which can produce strand breaks or mutations that alter transcription and DNA methylation, are primarily resolved by base excision repair (BER) (41). Importantly, we have demonstrated that genetic variants in the BER pathway confer PD risk in the context of environmental exposures (42). BER is initiated by lesion-specific glycosylases (e.g., OGG1) and completed by coordinated activities of APEX1, XRCC1, and DNA polymerase β (POLβ) (43,44). These BER proteins are critical for repairing DNA damage, specifically in promoters of actively transcribed neuronal genes (45–48). Poly (adenosine 5’-diphosphate-ribose) (PAR) polymerase-1 (PARP1) is a key DNA damage sensor for nicks, abasic sites, strand breaks, altered replication forks, and even R-loops (49–51). PARP1 is primarily involved in BER and single-strand break (SSB) repair (49). Upon recognition of DNA damage, PARP1 becomes activated and synthesizes poly-ADP-ribose (PAR) polymers derived from nicotinamide adenine dinucleotide (NAD^+^), on itself and other target proteins, which promotes the recruitment of downstream repair factors (49,52). The addition of ADP-ribose units alters the physical and enzymatic properties of acceptor proteins, which can influence the protein’s cellular activity and localization. PAR is rapidly removed by poly(ADP-ribose) glycohydrolase (PARG) to maintain PAR homeostasis (49,53). PAR can also serve as a scaffold for the recruitment of other DDR proteins to facilitate repair, such as XRCC1(49). PARP1 has well defined roles in the regulation of genome stability and has been linked to neurologic disorders (49,50,54–60).

Here, we examined the role of nuclear DNA damage and PARP-mediated signaling in *in vitro* and *in vivo* LRRK2 PD models. We found that LRRK2 G2019S causes endogenous nuclear oxidative DNA damage and PARP1-dependent PAR accumulation, with associated chromatin-retention of PARP1 and BER factors. PD-linked LRRK2 G2019S cells are selectively sensitive to DNA damaging agents that generate BER-substrate lesions as well as to PARP-trapping inhibitors, whereas a SOD/catalase mimetic rescues elevated PAR, thereby defining a ROS-driven PARP1 hyperactivation pathway that constitutes a targetable mechanism to leverage or mitigate nuclear DNA damage and downstream PARP1-mediated cytotoxicity.

## MATERIALS AND METHODS

### Mammalian cell culture and treatments

LRRK2^G2019S/G2019S^ KI HEK293 cells were generated using a CRISPR/Cas9 genome editing approach which was described previously (33). The LRRK2^G2019S/G2019S^ KI HEK293 and wild-type control cells were maintained in DMEM/F12, GlutaMAX™ supplement (Gibco, 10565-018), Seradigm Premium Grade 10% FBS (VWR, 97068-085) or HyClone™ Characterized Fetal Bovine Serum (Cytiva, SH30071.03), and 0.5% Penicillin Streptomycin (Corning, 30-002-Cl) in an incubator at 37 °C with 5% CO_2_. Plasmocin Prophylactic (InvivoGen) was added to media with cells at a concentration of 5 µg/ml.

HEK293 cells stably transfected with human LRRK2 or the G2019S variant of human LRRK2 (WT-LRRK2 and G2019S-LRRK2), were maintained in an incubator at 37 °C with 5% CO_2_ and grown in Eagle’s Minimum Essential Medium (ATCC: The Global Biosource, 30-2003), 10% Gibco Fetal Bovine Serum, Qualified (Fisher Scientific, 10-437-028) or HyClone™ Characterized Fetal Bovine Serum (Cytiva, SH30071.03), and 0.5% Penicillin-Streptomycin (Corning, 30-002-Cl). Geneticin (Thermo Fisher Scientific, 10131035) was added to media with cells at a concentration of 400 μg/ml.

### DNA damage treatments for cell viability assays

LRRK2^G2019S/G2019S^ KI HEK293 and wild-type cells were exposed to either H_2_O_2_, methyl methane sulfonate (MMS), cis-diammineplatinum(II) dichloride (cisplatin), ionizing radiation (IR), or hydroxyurea (HU). Cells were plated at a density of 0.4 × 10^6^ in a 6-well dish and treated at ∼40–50% confluency. Cells treated with H_2_O_2_ (Sigma Aldrich 216763) were incubated with 750 μM H_2_O_2_, 1 mM H_2_O_2_, or vehicle control for 1.5 hours (h). H_2_O_2_ was diluted in sterile-filtered water suitable for cell culture (Sigma-Aldrich, W3500-500ML). For MMS, cells were plated at a density of 0.4 × 10^6^ in a 6-well dish and incubated with 0.25 mg/mL MMS (Sigma Aldrich 129925), 0.5 mg/mL MMS, or vehicle control at ∼40–50% confluency for 1.5 h. MMS was diluted in complete growth medium.

Cisplatin-treated cells were plated at a density of 0.15 × 10^6^ in a 6-well dish and were subsequently incubated with 1 μM cisplatin (Sigma Aldrich P4394), 10 μM cisplatin, or vehicle control for 48 h. Cisplatin was resuspended and diluted in sterile-filtered water suitable for cell culture (Sigma-Aldrich, W3500-500ML). Cells that were exposed to IR were plated at a density of 0.2 × 10^6^ in a 6-well dish. They received 0, 5, or 10 Gy IR using the X-RAD 320 irradiator (Precision X-ray Inc., North Bradford, CT) and were stained 24 h after irradiation. Similarly, cells treated with HU (Selleck Chemicals S1896) were plated at a density of 0.15 × 10^6^ in a 6-well dish and then incubated with 200 μM HU, 400 μM HU, or vehicle control for 48 h. HU was resuspended and diluted in sterile-filtered water suitable for cell culture (Sigma-Aldrich, W3500-500ML). Cell viability was measured using annexin V/propidium iodide (PI) staining, as described below.

### PARP and PARG inhibitor treatments

To inhibit PARG activity, LRRK2^G2019S/G2019S^ KI HEK293 and wild-type cells were plated at a density of 0.4 × 10^6^ in a 6-well dish and treated at ∼40–50% confluency with 10 µM PARG inhibitor (Tocris, PPD 00017273) for 1.5 h. The PARG inhibitor was dissolved in DMSO (Sigma Aldrich, D2650-100ML). The same final DMSO concentration was used for each solution, which did not exceed 0.1% v/v. To inhibit PARP1/2 activity, LRRK2^G2019S/G2019S^ KI HEK293 and wild-type cells were treated with PARP1/2 inhibitors olaparib (APExBIO, A4154) or veliparib (Selleck Chemicals, S1004) at a concentration of 10 µM for 1.5 h. For viability experiments after PARP1/2 inhibition, LRRK2^G2019S/G2019S^ KI HEK293 and wild-type cells were treated with 2.5 and 5 µM Olaparib or 5 and 10 µM veliparib for 48 h. Olaparib and veliparib were dissolved in DMSO, and the same final DMSO concentration was used for each solution, which did not exceed 0.1% v/v.

To inhibit specific PARPs, LRRK2^G2019S/G2019S^ KI HEK293, WT-LRRK2, G2019S-LRRK2, and wild-type cells were plated at a density of 0.4 × 10^6^ in a 6-well dish and treated at ∼40–50% confluency. Cells were treated for 1.5 h with vehicle, 30 nM, or 300 nM of PARP1 inhibitor AZD5305 (Selleck Chemicals, S9875), vehicle, 3 µM, or 10 µM PARP2 inhibitor UPF 1069 (Selleck Chemicals, S8038), and vehicle, 3 µM, or 6 µM of PARP3 inhibitor ME0328 (Selleck Chemicals, S7438). PARP inhibitors (AZD5305, UPF 1069, and ME0328) were dissolved in DMSO. The same final DMSO concentration was used for each solution, which did not exceed 0.1% v/v.

### ROS scavenger treatments

LRRK2^G2019S/G2019S^ KI and wild-type HEK293 cells were plated at densities of 0.15 × 10^6^ and 0.2 × 10^6^ respectively, in 6-well dishes and treated for either 24 or 48 h, with vehicle, 20 µM, or 50 µM EUK-134 (SelleckChem, S4261). LRRK2^G2019S/G2019S^ KI and wild-type HEK293 cells were plated at a density of 0.15 × 10^6^ in 6-well dish and treated with vehicle, 0.5 µM, 5 µM, or 50 µM EUK-8 (Sigma Aldrich, 341209-10MG) for 48 h. EUK-134, and EUK-8 were dissolved and diluted in DMSO, and the same final DMSO concentration was used for each solution, which did not exceed 0.1% v/v.

### Rotenone and MMS treatments

LRRK2^G2019S/G2019S^ KI and wild-type HEK293 cells were plated at a density of 0.2 × 10^6^ in a 6-well dish and treated with vehicle, 10 nM, or 100 nM rotenone (Millipore Sigma, R8875-1G) for 24 h. Rotenone was dissolved and diluted in DMSO, and made fresh for each experiment. The same final DMSO concentration was used for each solution, which did not exceed 0.1% v/v. LRRK2^G2019S/G2019S^KI and wild-type cells were seeded at a density of 0.4 × 10^6^ in a 6-well dish, and treated with vehicle or 4 mM MMS for 25 min. For flow cytometry analysis of PAR, WT-LRRK2 and G2019S-LRRK2 cells were treated with vehicle or 0.5 µM MMS for 30 min.

### DRB treatments

LRRK2^G2019S/G2019S^ KI and wild-type HEK293 cells were plated at a density of 0.4 × 10^6^ in a 6-well dish and treated with vehicle, 20 µM, or 100 µM 5,6-Dichloro-1-β-D-ribofuranosylbenzimidazole (DRB) (Enzo, BML-EI231-0050) for 2 or 6 h. DRB was resuspended in DMSO. The same final DMSO concentration was used for each solution, which did not exceed 0.1% v/v.

### siRNA transfections

For siRNA knockdown experiments, LRRK2^G2019S/G2019S^ KI and wild-type HEK293 cells were plated at a density of 0.15 × 10^6^ in a 6-well dish. Cells were transfected with Dharmacon ON-TARGETplus SMARTPool siRNA for human PARP1 at 25 nM (Horizon Discovery, L-027012-00-0005), or a non-targeting siRNA (Horizon Discovery, D-001810-01-05) using 9 µL of Invitrogen Lipofectamine RNAiMAX transfection reagent per transfection (Fisher Scientific 13-778-150) in Opti-MEM Reduced Serum Media (Thermo Scientific 3195070). The following day, cell media was replaced with complete growth medium and cells were allowed to recover for 24 h. Knock-down efficiency of PARP1 was validated by quantitative western blot.

### Annexin V/PI staining and flow cytometry

Cell viability was measured using annexin V/propidium iodide (PI) staining (ThermoFisher Scientific, V13245) and flow cytometry. Annexin V and PI staining was performed according to the manufacturer’s instructions. Cells were incubated with pharmacological inhibitors or siRNAs and transfection reagents, as described above. Media with detached cells was collected and combined with adhered cells after retrieval with trypsin-EDTA (ThermoFisher Scientific, 25300062) or versene solution (ThermoFisher Scientific, 15040066). Cells were centrifuged at 500 × g for 5 minutes, washed with cold PBS, and centrifuged at 500 × g for 3 minutes. Cells were diluted in 1X annexin-binding buffer to ∼1 × 10^6^ cells/mL. 5 µL Alexa Fluor 488 Annexin V and 1 µL 100 µg/mL PI was added to each 100 µL of cell suspension and incubated at room temperature for 15 minutes. After the incubation period, 400 µL 1X annexin-binding buffer was added to each sample, mixed gently, and kept on ice until analysis. Cells were analyzed on a BD FACSCanto-II flow cytometer (BD Biosciences). BD FACS Diva software 6.0 (BD Biosciences) and FlowJo software (BD Biosciences) were used for the gating of live (Annexin V/PI double negative), early apoptotic (Annexin V-positive, PI-negative), late apoptotic (Annexin V/PI double positive, and dead (Annexin V-negative/PI-positive) cells. Ten thousand cells gated as single cells were analyzed.

### Oxidative Repair Assisted Damage Detection (oxRADD)

LRRK2^G2019S/G2019S^ KI and wild-type control HEK293 cells were plated on coverslips (Electron Microscopy Sciences, 72290-04) coated with poly-D-lysine (Sigma Aldrich, P6407) and fixed using 4% paraformaldehyde (Electron Microscopy Sciences, 15710) with MgCl_2_ (Sigma Aldrich, M2392) in PBS for 10 minutes at room temperature. They were then washed three times with PBS and stored at 4℃ until processed. Cells were permeabilized using permeabilization buffer (Biotium, Cat. # 22016) with 0.05% Triton X-100 (Sigma Aldrich, X100-5ml) for 10 minutes at room temperature, then washed twice with PBS. For lesion removal, cells were washed once with water, and then incubated with 100 µL of a lesion removal mix consisting of 10 µL Thermo Pol Buffer (New England Biolabs, B9004), 500 µM NAD^+^ (New England Biolabs, B9007), 200 µg/mL BSA (Sigma Aldrich, A2058), 4 U FPG (NEB, M0240), 5 U EndoIV (New England Biolabs M0304), and 5 U EndoVIII (New England Biolabs, M0299) for 1 h at 37℃. Cells were then incubated with 100 µL of a gap-filling mix consisting of 1 µL Klenow exo− (ThermoFisher, EP0422), 0.1 µL Digoxigenin dUTP (Sigma Aldrich, 11093088910), 200 µg/mL BSA, and 10 µL Thermo Pol Buffer (New England Biolabs, B9004) for 1 h at 37℃. Cells were washed three times with PBS, and blocked with 5% goat serum in PBS for 30 minutes at room temperature. Anti-Digoxigenin antibody (Abcam ab420, 1:250) was diluted in 5% goat serum in PBS and incubated at room temperature for 1 h. Cells were subsequently washed three times with PBS and incubated with Mouse Alexa Fluor 546 secondary antibody (Life Tech) diluted in 5% goat serum in PBS for 1 h at room temperature. Cells were then washed three times with PBS and incubated with Hoechst (1:1000) for 15 minutes at room temperature. Cells were washed once more three times with PBS and mounted with Prolong Gold. For isotype control experiments to confirm specificity, monoclonal mouse IgG1 (Cell Signaling 5415) was used at an equivalent concentration to the anti-Digoxigenin antibody on cells and stained as described. After 24 h, samples were stored at 4℃ until imaging.

Cells were then imaged using the Keyence microscope with the 20× objective (NA 0.75). At least 100 cells from each condition and biological replicate were collected across 5 different fields. For analysis, the Nikon Elements software created an ROI around the nucleus, and the total intensity for the RADD channel within the nucleus was recorded. The fluorescent intensity for the nuclei was graphed in GraphPad Prism, and significance was calculated using an unpaired t-test for each DNA adduct category versus its matched control.

### Western blot analysis for cell lines

Whole cell lysates were collected using lysis buffer consisting of RIPA buffer, protease inhibitor cocktail (Sigma Aldrich, P8340), and Halt phosphatase inhibitor (ThermoFisher Scientific, 78420). After incubating on ice for 10 minutes, cell lysates were centrifuged at 10,000 × g for 15 minutes at 4°C. Cell lysates were stored at -80°C until used for immunoblotting. Protein was quantified with the DC protein assay (Bio-Rad, 5000116) according to the manufacturer’s instruction manual. For western blots, cell lysates were incubated at 100°C for 5 minutes with NuPAGE sample loading dye (ThermoFisher Scientific, NP0007) and dithiothreitol (Bio-Rad 1610610). Protein lysates were run on 4-20% mini-protean TGX stain-free gels (Bio-Rad, 4568096) and transferred using the Trans-Blot Turbo Transfer System (Bio-Rad 1704155) and Trans-Blot Turbo RTA Midi 0.2µm Nitrocellulose Transfer Kit (Bio-Rad, 1704271). Membranes were blocked in 5% w/v nonfat dry milk in PBST (PBS with 0.05% Tween 20) and incubated with primary antibodies/binding reagents diluted in 5% w/v nonfat dry milk in PBST at 4°C overnight (anti-poly-ADP-ribose binding reagent, Millipore Sigma, MABE1031, 1:10,000; anti-PARP1, Santa Cruz Biotechnology, sc-8007, 1:1,000; anti-XRCC1, Novus Biological, NBP1-87154, 1:5,000; anti-Pol β, Abcam, ab26343, 1:1,000; anti-Ligase III, GeneTex, GTX70143, 1:1,000; anti-H3, Abcam, ab1791, 1:20,000; anti-ɑ-tubulin, Abcam, ab7291, 1:10,000; anti-β-actin, Novus Biologicals, NBP1-47423, 1:10,000; anti-β-actin, Abcam, ab8227, 1:10,000). Membranes were incubated with LI-COR IRDye secondary antibodies (IRDye 680R donkey anti-mouse, LI-COR, 926-68072, 1:10,000; IRDye 680RD donkey anti-rabbit, LI-COR, 926-68073, 1:10,000; IRDye 800CW donkey anti-mouse, LI-COR, 926-32212, 1:10,000; IRDye 800CW donkey anti-rabbit, LI-COR, 926-32213, 1:10,000) diluted in 5% w/v nonfat dry milk in PBST for 1 h at room temperature and scanned using an Odyssey M Imager (LI-COR). Quantification was performed with Image Studio Lite (LI-COR) software.

### Mice

All animals were used in accordance with the Institutional Animal Care and Use Committee (IACUC) and the Duke Division of Laboratory Animal Resources (DLAR) oversight (IACUC Protocol Number A036-22-02-25). All mice used were 4-6 months of age and included male and females. Non-transgenic C57BL/6J wild-type (WT) control mice and the *Lrrk2* G2019S knock-in heterozygotes or homozygote (GKI) mice, developed in the laboratory of Heather Melrose (61), were maintained at Duke University School of Medicine. The resulting offspring were genotyped by tail biopsy with previously described primers (61). This mouse strain is commercially available (Jackson Laboratory, #030961). Anesthetized animals were transcardially perfused with saline and decapitated. Brains were removed, rinsed in PBS, and microdissected on ice and then flash-frozen in liquid nitrogen and stored at -80°C until processed for immunoblotting.

### Western blot analysis for mouse brain tissue

Microdissected brain samples were homogenized with a Dounce tissue homogenizer in lysis buffer. After incubating on ice for 10 minutes, homogenized samples were centrifuged at 16,000 ×g for 15 minutes at 4°C. Supernatant was collected and stored at -80°C until protein was quantified. Protein was quantified with the DC protein assay (Bio-Rad, 5000116) according to the manufacturer’s instruction manual. For western blots, whole cell brain lysates were incubated at 70°C for 5 minutes with NuPAGE sample loading dye (ThermoFisher Scientific, NP0007) and dithiothreitol (DTT) (Bio-Rad, 1610610). Protein lysates were run on 4-20% mini-protean TGX stain-free gels (Bio-Rad, 4568096) and transferred using the Trans-Blot Turbo Transfer System (Bio-Rad, 1704155) and Trans-Blot Turbo RTA Midi 0.2µm Nitrocellulose Transfer Kit (Bio-Rad, 1704271). Membranes were blocked in 5% w/v nonfat dry milk in PBST (PBS with 0.05% Tween 20) and incubated with primary antibodies/binding reagents diluted in 5% w/v nonfat dry milk in PBST at 4°C overnight (anti-poly-ADP-ribose binding reagent, Millipore Sigma, MABE1031, 1:10,000; Novus Biologicals, NBP1-47423, 1:10,000). Membranes were incubated with LI-COR IRDye secondary antibodies (IRDye 680R donkey anti-mouse, LI-COR, 926-68072, 1:10,000; IRDye 800CW donkey anti-rabbit, LI-COR, 926-32213, 1:10,000) diluted in 5% w/v nonfat dry milk in PBST for 1 h at room temperature and scanned using an Odyssey M Imager (LI-COR). Quantification was performed with Empiria Studio (LI-COR) software.

### Immunofluorescence

For imaging experiments, LRRK2^G2019S/G2019S^ KI and wild-type control HEK293 cells were plated on coverslips (Electron Microscopy Sciences, 72290-04) coated with poly-D-lysine (Sigma Aldrich, P6407) and fixed using 4% paraformaldehyde (Electron Microscopy Sciences, 15710) with MgCl_2_ (Sigma Aldrich, M2392) in PBS for 10 minutes at room temperature. Cells were permeabilized with 0.5% Triton X-100 (Sigma Aldrich, T8787) in PBS for 10 minutes, then blocked with 3% BSA in PBS for 30 minutes. After blocking, cells were incubated in primary antibodies (anti-poly-ADP-ribose binding reagent, Millipore Sigma, MABE1031, 1:2,000; anti-PARP1, Proteintech, 66520-1-Ig, 1:500) diluted in 3% BSA in PBS overnight at 4°C. Cells were then washed with PBS and incubated with Invitrogen Alexa Fluor donkey secondary antibodies (Donkey anti-Rabbit IgG (H+L) Secondary Antibody, Alexa Fluor 488, ThermoFisher Scientific, A21206, 1:500; Donkey anti-Mouse IgG (H+L) Highly Cross-Adsorbed Secondary Antibody, Alexa Fluor 488, ThermoFisher Scientific, A21202, 1:500) for 1 h. Following incubation with secondary antibodies, cells were washed with PBS and stained with Hoechst (Invitrogen ThermoFisher Scientific, H3570). Finally, slides were rinsed with H_2_O and mounted with Prolong Diamond mounting media (ThermoFisher Scientific, P36970). To analyze immunofluorescence, slides were imaged on a Zeiss 780 upright confocal microscope with a 63x oil immersion objective. 405 nm Diode, 488 nm Argon/2, and 561 nm Diode lasers were used for imaging. QuPath (version 0.5.1) was used for image analysis and quantification.

### Mitochondrial Replication Assay (MIRA)

To quantify cell populations in S-phase and analyze EdU incorporation, we utilized the Mitochondrial Replication Assay (MIRA) protocol (62,63). WT-LRRK2 or G2019S-LRRK2 cells were plated on 8-well chamber microscope slides (Thermo Scientific, 177402) at 35,000 cells/well and were allowed to grow for 24 h at 37 °C, 5% CO_2_. After 24 h of incubation, media was aspirated and replaced with media containing 20 μM 5′-ethylene-2′-deoxyuridine (EdU) (Invitrogen, A10044), and cultures were allowed to incubate for 1 h. After EdU treatment, cells were washed two times with PBS (Thermo Scientific, 10010049), and fixed with 2% paraformaldehyde (Electron Microscopy Sciences, 15710) in PBS for 15 minutes at room temperature. Following fixation, cells were washed twice with PBS for 5 minutes each with shaking at ∼25 rpm at room temperature, and cell culture chambers and silicone gasket were removed from the chamber slides. Cells were permeabilized by placing slides in a Coplin jar containing 60 mL of 0.25% Triton X-100 (Sigma-Aldrich, T8787-100ML) in PBS with slow shaking at room temperature for 15 minutes, followed by three 5 minute washes in a Coplin jar filled with PBS. A Click-IT chemical reaction was then used to label EdU incorporated into cellular DNA. The Click-IT reaction cocktail was made by adding 2 mM copper sulfate (Fluka Analytical, 35185), 10 μM biotin azide (Invitrogen, B10184) + Alexa 488-azide (Invitrogen, A10266), and 100 mM sodium ascorbate (Sigma-Aldrich, 11140) to PBS in that order, with vortexing following addition of each constituent. A humidified slide chamber was created by placing wet paper towels inside of a slide box (VWR, 82003-414), and chamber slides were placed face-up in the humidified chamber. 45 μL of the Click-IT reaction cocktail was added to each well of the chamber slides, the slides were covered with plastic coverslips (Electron Microscopy Sciences, 72261-50), the humidified chamber was closed, and the slides were allowed to incubate at room temperature for 1 h. After incubation, plastic coverslips were removed and the slides were washed in PBS at room temperature for 5 minutes three times. The slides were then placed back into the humidified chamber, and 50 μL of Duolink blocking solution (Sigma-Aldrich, DUO82007-8ML) was added to each chamber slide well. Slides were covered with new plastic coverslips, the humidified chamber was closed, and slides were incubated with blocking solution at room temperature for 1 h. Coverslips were then removed, and 35 μL of mouse anti-biotin (1:100, Sigma-Aldrich, 7653) and rabbit anti-biotin (1:100, Cell Signaling, 5597S) diluted in Duolink antibody diluent (Sigma-Aldrich, DUO82008-8ML) was added to each well. Slides were covered with new coverslips and incubated in the humidified chamber at 4°C overnight. A proximity ligation assay was performed following incubation with primary antibodies. The slides were washed three times for 5 minutes in a Coplin jar with wash buffer A, which consisted of 10 mM Trizma hydrochloride (Sigma-Aldrich, T5941), 150 mM sodium chloride (Sigma-Aldrich, S7653-250G), and 0.05% by volume Tween 20 (Fisher Scientific, BP337-100) in autoclaved Milli-Q water, pH adjusted to 7.5 using hydrochloric acid (Sigma-Aldrich, 1090631000). The slides were then placed in the humidified chamber, pre-warmed to 37°C, and 30 μL Duolink antibody diluent containing 1:5 Duo link in situ anti-mouse (Sigma-Aldrich, DUO92001-100RXN) and anti-rabbit (Sigma-Aldrich, DUO92005-100RXN) proximity ligation assay (PLA) probes were added to each well. Slides were covered with clean plastic coverslips and incubated in the humidified chamber at 37°C for 1 h. After incubation, slides underwent three 5 minute washes in wash buffer A. Duolink in situ detection reagents (Sigma-Aldrich, DUO92008-100RXN) were used to perform a PLA reaction to enhance detection of EdU nucleotide incorporation. The Duolink ligase (1:40) and Duolink ligation buffer (1:5) were diluted in autoclaved Milli-Q water, and 30 μL were added to each slide well. A clean plastic coverslip was added, and the slides were incubated at 37°C in the humidified chamber for 30 additional minutes, followed by two 5-minute washes in wash buffer A. The amplification mix was prepared by adding the Duolink polymerase (1:80) and amplification buffer (1:5) in autoclaved Milli-Q water, and 30 μL per well were added to the slides, which were covered and incubated in the humidified chamber at 37°C for 100 minutes. After amplification, the slides were washed three times for 10 minutes at room temperature in wash buffer B, which consisted of 200 mM Trizma hydrochloride and 100 mM sodium chloride in autoclaved Milli-Q water, pH adjusted to 7.5. These washes occurred in a Coplin jar covered with tinfoil to protect slides from light. Finally, the slides were washed in 0.01x wash buffer B for 1 minute in a covered Coplin jar. Nuclei were stained by adding 30 μL 1:1000 Hoescht 33342 solution (Thermo Fisher Scientific, H3570) in PBS to each well, covering slides with fresh coverslips, and incubating in the humidified chamber for 5 minutes at room temperature. Slides were then washed two times in PBS at room temperature for 5 minutes, followed by mounting using Prolong Gold antifade reagent (Invitrogen, P36934) and glass coverslips (Fisher Scientific, 12-548-5M). Mounted slides were stored overnight at room temperature in the dark and immediately imaged the next day. The analysis was derived from three independent biological replicates. The following controls were utilized to ensure PLA signal specificity; these included separate conditions with no EdU treatment, no Click-IT reaction, no primary antibody (mouse and/or rabbit), no Duolink PLA probes (mouse and/or rabbit), no amplification mix, and no ligation mix. Slides were imaged using 405 nm Diode, Argon/2 488 nm, and 561 nm Diode lasers with a 63x oil immersion objective on a Zeiss 780 upright confocal workstation. For analysis, images were analyzed in QuPath (v0.5.1) to quantify average nuclear PLA puncta per cell and percent of Alexa 488-azide-positive or negative cells.

### Subcellular fractionation and quantification of soluble nuclear and chromatin-bound factors

Subcellular protein fractionations were performed using the Subcellular Protein Fractionation Kit for Cultured Cells (ThermoFisher Scientific, 78840). LRRK2^G2019S/G2019S^ KI and wild-type HEK293 cells were plated at a density of 0.5 × 10^6^ in a 6-well dish. Three wells were used for each condition. The following day, cells were harvested with trypsin-EDTA and centrifuged at 500 × g for 5 minutes. Cells were resuspended in ice-cold PBS. Approximately 2 × 10^6^ cells were centrifuged at 500 × g for 3 minutes. Cell pellets were resuspended in 100 µL ice-cold cytoplasmic extraction buffer containing protease inhibitors and incubated on a revolver rotator at 4°C for 10 minutes. Lysates were centrifuged at 500 × g for 5 minutes and supernatants were transferred to a pre-chilled tube on ice. 100 µL ice-cold membrane extraction buffer containing protease inhibitors was added to the pellet and vortexed for 5 seconds and incubated on a rotator at 4°C for 10 minutes. Lysates were centrifuged at 3,000 × g for 5 minutes and supernatants were transferred to a pre-chilled tube on ice. 50 µL ice-cold nuclear extraction buffer containing protease inhibitors was added to the pellet and vortexed for 15 seconds and incubated on a rotator at 4°C for 10 minutes. Lysates were centrifuged at 5,000 × g for 5 minutes and supernatants were transferred to a pre-chilled tube on ice. Finally, 50 µL room temperature nuclear extraction buffer containing protease inhibitors, CaCl_2_, and Micrococcal Nuclease was added to the pellet and vortexed for 15 seconds. Samples were incubated at 37°C for 5 minutes and centrifuged at 16,000 × g for 5 minutes and supernatants were transferred to a pre-chilled tube on ice. All centrifugation steps were performed at 4°C. Lysates were stored at -80°C until protein was quantified. Protein was quantified with the DC protein assay (Bio-Rad, 5000116) according to the manufacturer’s instruction manual.

### RNA purification and quantitative real-time PCR (qRT-PCR)

Cells were plated at a density of 0.3 × 10^6^ in a 6-well dish for 24 hours. Three wells were collected for each genotype, achieving a final total of ∼ 1 x10^6^ cells. Total cellular RNA was isolated from wild-type control and LRRK2^G2019S/G2019S^ KI HEK293 cells using the RNeasy kit (Qiagen, 74104) according to the manufacturer’s protocol and stored at -80°C until conversion to cDNA. cDNA was generated with the RT^2^ First Strand Kit (Qiagen, 330404). After cDNA conversion, qRT-PCR was performed using primers for PARP1 (Qiagen PPH00686B and PPM05150C) and XRCC1 (Qiagen PPH01741A and PPM04396A). qRT-PCR reactions consisted of PowerUp SYBR Green Mastermix (Applied Biosystems A25777P), primers, and nuclease-free water (Qiagen, 129114). Amplification and measurement were completed using a QuantStudio 3 Real-Time PCR System (Applied Biosystems). Amplicon product specificity was confirmed by melt curve analysis and gel electrophoresis.

### PAR quantification by flow cytometry

WT-LRRK2 and G2019S-LRRK2 cells were plated at a density of 0.5 × 10^6^ in a 6-well dish. 24 h after seeding, cells were treated with vehicle or 0.5 μM MMS for 0.5 h. Cells were collected with 0.25% Trypsin-EDTA (Thermo Fisher Scientific, 25200072) and washed with PBS. 1 × 10⁶ cells per sample were transferred to round bottom tubes (Corning, 352058). 1 mL of FACS buffer was added to each tube, and cells were stained with Zombie Yellow Fixable Viability Dye (Biolegend, 423103). Cells were then fixed and permeabilized using the Foxp3/Transcription Factor Staining Buffer Set (Fisher Scientific, 00-5523-00) according to the manufacturer’s instructions. Briefly, cells were incubated in 200 μL fixation/permeabilization solution (diluted 1:3) for approximately 45 minutes at room temperature, followed by two washes with permeabilization buffer. Cells were subsequently incubated with anti-PAR/pADPr primary antibody (clone 10H) (Bio-Techne Sales Corp., Catalog # NBP2-89039) diluted in permeabilization buffer. After washing, cells were incubated with a PE-conjugated secondary antibody (Bio-Techne Sales Corp., F0102B, 10 µL per 1 × 10⁶ cells) for 30 minutes at room temperature, protected from light. Following two additional washes with permeabilization buffer, cells were resuspended in FACS buffer and kept on ice until acquisition. Flow cytometry was performed on a BD FACSCanto-II cytometer, and the percentage of PAR-positive cells was quantified using FlowJo software.

### Statistical Analyses

Data were analyzed with GraphPad Prism. Outliers were determined with Grubb’s test. Statistical significance was defined by p<0.05. For all graphs, values and error bars are presented as mean ± standard error of mean (SEM).

## RESULTS

### Enhanced sensitivity to DNA-damaging agents in PD-linked mutant LRRK2 G2019S cells

The PD-linked variant LRRK2 G2019S has been implicated in cellular stress responses, but its effects on DNA damage tolerance and repair pathways remain unclear. To address this, we compared the viability of CRISPR/Cas9-generated homozygous knock-in (LRRK2^G2019S/G2019S^ KI) with wild-type HEK293 control cells following exposure to multiple genotoxic agents with distinct mechanisms of action to determine whether LRRK2 G2019S causes broad defects in damage tolerance/repair or is potentially lesion or pathway specific. Cell death was quantified by Annexin V/propidium iodide (PI) staining and flow cytometry (Figure 1A). All treatment responses are reported relative to each genotype’s baseline viability to account for the lower baseline survival observed in LRRK2^G2019S/G2019S^ KI cells compared to wild-type (Supplemental Figure 1).

**Figure 1.**
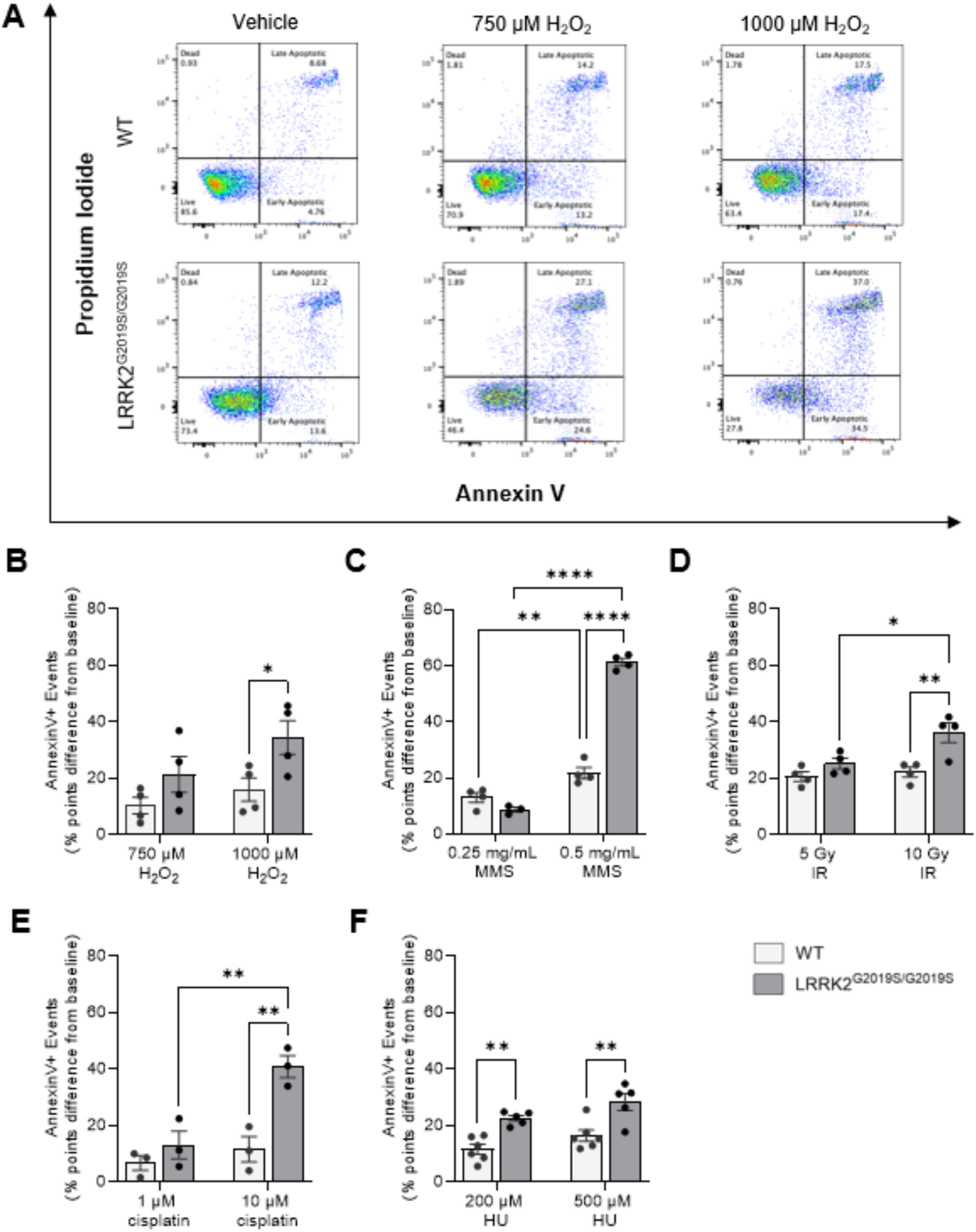
LRRK2^G2019S/G2019S^ KI cells exhibit apoptosis sensitivity following treatment with genotoxic insults compared to wild-type cells. Wild-type (WT) and LRRK2^G2019S/G2019S^ KI cells were exposed to (A, B) hydrogen peroxide (H_2_O_2_), (C) methyl methanesulfonate (MMS), (D) ionizing radiation (IR), (E) cisplatin, or (F) hydroxyurea (HU) and stained for annexin V/propidium iodide (PI). (A) Representative flow cytometry plots of live, apoptotic, and necrotic cell populations in wild-type and LRRK2^G2019S/G2019S^ KI cells treated with vehicle or H_2_O_2_. (B) Analysis of Annexin V-positive cells following treatment with vehicle, 750 μM or 1 mM H_2_O_2_ for 1.5 h (n=4). (C) Flow cytometry analysis of apoptotic cells after incubation with vehicle, 0.25 mg/mL or 0.5 mg/mL MMS for 1.5 h (n=4). (D) Flow cytometry analysis of cells exposed to 0, 5, or 10 Gy IR and stained 24 h later (n=4). (E) Flow cytometry analysis of cells treated with vehicle, 1 μM or 10 μM cisplatin for 48 h (n=3). (F) Flow cytometry analysis of cells incubated with vehicle, 200 μM HU or 500 μM HU for 48 h (n=6). Values are presented as percent points difference from either the WT or LRRK2^G2019S/G2019S^ KI cell baseline. *p < 0.05, **p < 0.01, ****p < 0.0001, as determined by two-way ANOVA with Bonferroni’s multiple comparison. n = 3 to 6 biological replicates. Data are presented as mean ± SEM.

To investigate the effect of an oxidizing agent, LRRK2^G2019S/G2019S^ KI and wild-type control cells were treated with multiple concentrations of hydrogen peroxide (H_2_O_2_) or vehicle. Acute exposure to H_2_O_2_ produced an increase in apoptosis in both genotypes, but LRRK2^G2019S/G2019S^ KI cells exhibited significantly greater apoptosis than wild-type at the higher concentration tested (1000 µM) (Figure 1A, B). Since H_2_O_2_ produces ROS and oxidative DNA base damage primarily repaired by BER, these results suggested the PD LRRK2 mutant increased sensitivity to oxidative lesions and potentially reduced BER capacity. We therefore next tested the sensitivity of LRRK2^G2019S/G2019S^ KI and wild-type control cells to the alkylating agent methyl methanesulfonate (MMS), since lesions generated by MMS are also primarily repaired by BER. LRRK2^G2019S/G2019S^ KI cells exposed to MMS induced dose-dependent cell death in both lines, with LRRK2^G2019S/G2019S^ KI cells showing a significant decrease in viability at 0.5 mg/mL MMS (Figure 1C, Supplemental Figure 2A).

To probe lesion-specific repair capacity, in particular DNA strand-breaks, cells were exposed to a high dose of ionizing radiation (IR, 5Gy), which reduced survival independent of genotype (Figure 1D). However, at 10Gy, a dose that produces extensive apoptosis/necrosis and confounds repair specificity, LRRK2^G2019S/G2019S^ KI cells showed a significant decrease in survival relative to wild-type (Figure 1D, Supplemental Figure 2B). We then probed the sensitivity of tolerance of crosslink repair. Exposure to the DNA crosslinking agent cisplatin for 48 hours (h) produced genotype-dependent differences only at the higher dose: 10 μM cisplatin reduced viability to a greater extent in LRRK2^G2019S/G2019S^ KI cells than in wild-type (Figure 1E, Supplemental Figure 2C). Though BER is not the primary pathway to repair DNA damage induced by cisplatin and low doses of IR, BER can serve as a backup or as a secondary repair pathway (64).

To test whether cells are hypersensitive to replication stress, cells were treated with hydroxyurea (HU), which inhibits ribonucleotide reductase, depleting the dNTP pools, stalling replication forks and promoting replication-associated single-strand breaks and fork collapse (65). HU exposure to induce replication stress resulted in increased apoptosis in a concentration-dependent manner in both cell lines. At both 200 μM and 500 μM HU, LRRK2^G2019S/G2019S^ KI cells displayed significantly higher Annexin V/PI positivity than wild-type controls (Figure 1F, Supplemental Figure 2D). Together, these results indicate that LRRK2 G2019S sensitizes cells to diverse genotoxic insults, consistent with impairment of one or more shared damage-sensing, signaling, or repair processes, including a preferential combination of those that resolve oxidative lesions and alkylation damage repaired by BER.

### LRRK2 G2019S drives endogenous oxidative DNA damage and elevated PAR in cells and mouse brains

Given the differential lethality of LRRK2^G2019S/G2019S^ KI cells to oxidative and alkylating agents, potentially reflecting an impaired ability to process these types of DNA lesions, we next assessed whether LRRK2 G2019S promotes accumulation of nuclear DNA damage under endogenous conditions. To do this, we quantified the presence of oxidative DNA lesions using the oxidative Repair Assisted Damage Detection (oxRADD) assay [53, 54]. In this assay, an enzyme cocktail containing oxidative lesion-specific DNA repair enzymes (FPG, EndoIV, and EndoVIII) converts oxidative base damage into gapped DNA and strand breaks, which are then filled with a labeled deoxyuridine triphosphate (dUTP) to enable fluorescent detection of oxidative DNA lesion sites within the genomic DNA. Using oxRADD, we detected low basal levels of oxidative DNA damage in wild-type cells (Figure 2A, B). In contrast, LRRK2^G2019S/G2019S^ KI cells showed a significant increase in endogenous oxidative DNA lesions by oxRADD compared with wild-type controls (Figure 2A, B). Only minimal fluorescent signal was detected in the isotype controls for either cell line (Supplemental Figure 3A, B).

**Figure 2.**
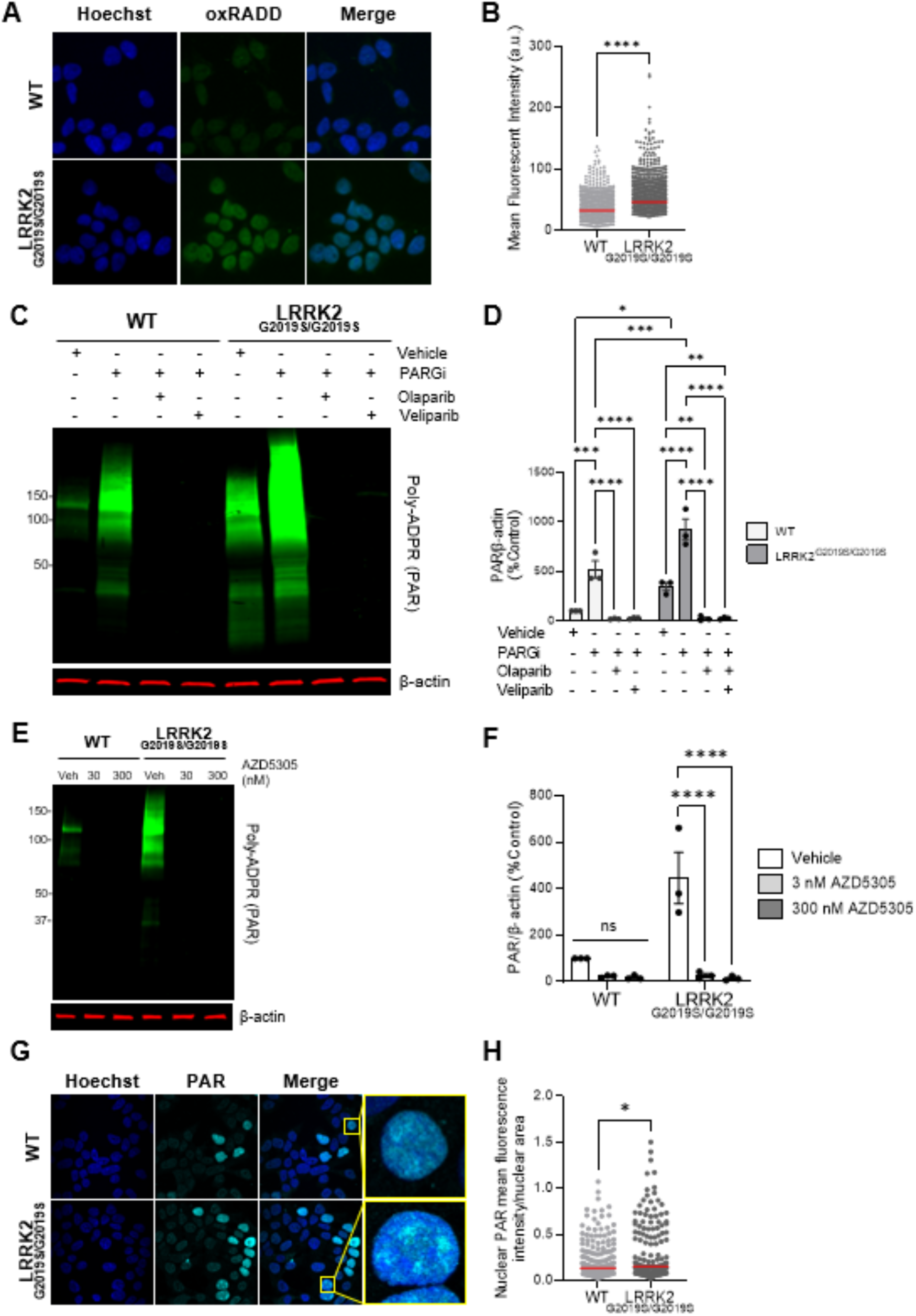
LRRK2 G2019S causes oxidative DNA damage and PARP1 activation *in vitro*. (**A**) Representative 20X wide-field images of wild-type (WT) and LRRK2^G2019S/G2019S^ KI cells incubated with Hoechst (blue) and oxRADD cocktail (green). (**B**) Quantification of oxidative lesions detected via the oxRADD assay. The red bar indicates the mean and each point represents an individual cell. Approximately 3500 total cells were analyzed from three independent experiments. (n=3; ****p < 0.0001, determined by unpaired t-test). (**C**) Representative western blot of wild-type and LRRK2^G2019S/G2019S^ KI cells treated with vehicle, a PARG inhibitor (10 mM), or a PARP1/2 inhibitor (10 mM, olaparib, veliparib) for 1 h and assessed for PAR levels and β-actin as a loading control. (**D**) Quantification of PAR levels showing elevation in LRRK2^G2019S/G2019S^ KI cells at baseline and following PARG inhibition compared to control cells. Olaparib and veliparib completely abolished PAR signal in both cell lines. (n=3, *p < 0.05, **p < 0.01, ***p < 0.001, ****p < 0.0001, as determined by two-way ANOVA with Bonferroni’s multiple comparison). (**E**) Representative western blot of wild-type and LRRK2^G2019S/G2019S^ KI cells treated with vehicle or the PARP1-selective inhibitor AZD 5305 (30 and 300 nM) for 1 h and assessed for PAR levels and β-actin as a loading control. (**F**) Quantification of PAR levels demonstrates PARP1-dependence in cells regardless of genotype. (n=3, ****p < 0.0001, determined by two-way ANOVA with Bonferroni’s multiple comparison). (**G**) Representative 60 x immunofluorescence confocal images and insets of wild-type and LRRK2^G2019S/G2019S^ KI cells incubated with Hoechst (blue) and PAR (cyan). (**H**) Quantification of nuclear PAR fluorescence intensity normalized to nuclear area revealed increased nuclear PAR signal in LRRK2^G2019S/G2019S^ KI cells relative to wild-type. The red bar indicates the mean and each point represents an individual cell. Data are from four independent experiments in which 120–160 nuclei were counted for each experiment. (n=4, *p < 0.05, determined by unpaired t-test). Data are presented as mean ± SEM.

The endogenous elevated oxidative DNA damage in LRRK2^G2019S/G2019S^ KI cells detected by oxRADD are lesions primarily repaired by BER and single-strand break (SSB) repair (SSBR), which activate poly (adenosine 5’-diphosphate-ribose) (PAR) polymerase-1 (PARP1). Therefore, we next examined PARP1 signaling, a key DNA damage sensor that responds to a wide range of lesions, including nicks, abasic sites, and strand breaks, and upon binding synthesizes PAR and initiates assembly of BER/SSBR complexes (49,50). Under endogenous conditions, LRRK2 G2019S caused a nearly 250 percent increase in PAR levels compared to the wild-type control using quantitative western blotting (Figure 2C, D). Since PAR polymers are rapidly catabolized by enzymes such as PARG, LRRK2^G2019S/G2019S^ KI and wild-type cells were also treated with a PARG inhibitor for 1 h. Following PARG inhibition, PAR levels increased approximately five-fold in wild-type cells (Figure 2C, D). Interestingly, with PARG inhibition, we observed a nearly two-fold increase in PAR accumulation in LRRK2^G2019S/G2019S^ KI cells compared to the wild-type control under similar conditions (Figure 2C, D).

To determine whether the LRRK2 G2019S-induced PAR generation is PARP1, PARP2 and/or PARP3 dependent, we first treated LRRK2^G2019S/G2019S^ KI and wild-type cells with the PARP1/2 inhibitors olaparib and veliparib. Treatment with PARP1/2 inhibitors olaparib or veliparib completely abolished LRRK2-mediated and wild-type derived PAR signal (Figure 2C, D). To probe PARP selectivity, we next acutely treated with the PARP1-selective inhibitor AZD5305, which fully abrogated PAR accumulation in both LRRK2^G2019S/G2019S^ KI and wild-type cells (Figure 2E, F). In contrast, exposure to the PARP2-selective inhibitor UPF1069, even at concentrations well above the IC_50_, did not alter PAR levels in either LRRK2^G2019S/G2019S^ KI or wild-type cells (Supplementary Figure 4 A, B). Treatment with the PARP-3 selective inhibitor ME0238 also had no effect on PAR levels in either genotype (Supplementary Figure 4 C, D). Together, these data demonstrate that the elevated PAR levels caused by LRRK2 G2019S are mainly PARP1 dependent.

To further investigate the LRRK2 G2019S generated elevated PAR levels, we next examined PAR localization by immunocytochemistry in LRRK2^G2019S/G2019S^ KI and wild-type control cells (Figure 2G). In both genotypes, PAR staining is largely co-localized with Hoechst, with non-detectable PAR signal in the cytoplasm by this method, indicating that basal levels of PAR are primarily generated in the nucleus (Figure 2G). Consistent with the quantitative PAR western blots (Figure 2C, D), nuclear PAR levels were increased in LRRK2^G2019S/G2019S^ KI cells relative to the wild-type control and inhibition of PARP1/2 with olaparib completely abrogated PAR signal (Figure 2G, H, Supplementary Figure 5).

We next compared PAR levels by quantitative western blotting in HEK293 cells stably transfected with human wild-type LRRK2 or the G2019S variant of human LRRK2 (hereafter WT-LRRK2 and G2019S-LRRK2, respectively) (27). Despite similar LRRK2 expression between the two cell lines (27), G2019S-LRRK2 cells exhibited a nearly four-fold increase in PAR levels relative to WT-LRRK2 expressing cells, suggesting a mutation-associated gain-of-function effect (Figure 3A, B). The G2019S-LRRK2 dependent PAR increase was not due to differences in the number of cells in S-phase, as quantified by Alexa 488 in the Mitochondrial Replication Assay (MIRA) assay, which employs EdU labeling of both nuclear and mitochondrial DNA and Click-IT chemistry to label EdU-tagged DNA with Alexa 488 azide and biotin azide (Figure 3C, D) (62). Interestingly the nuclear MIRA signal, obtained through PLA labeling of biotin-tagged EdU incorporated into nuclear DNA, was increased both in cells in and outside of S-phase in the G2019S-LRRK2 relative to the WT-LRRK2 cells; no signal was detected with appropriate PLA controls (Figure 3E,F, Supplemental Figure 6A, B).

**Figure 3.**
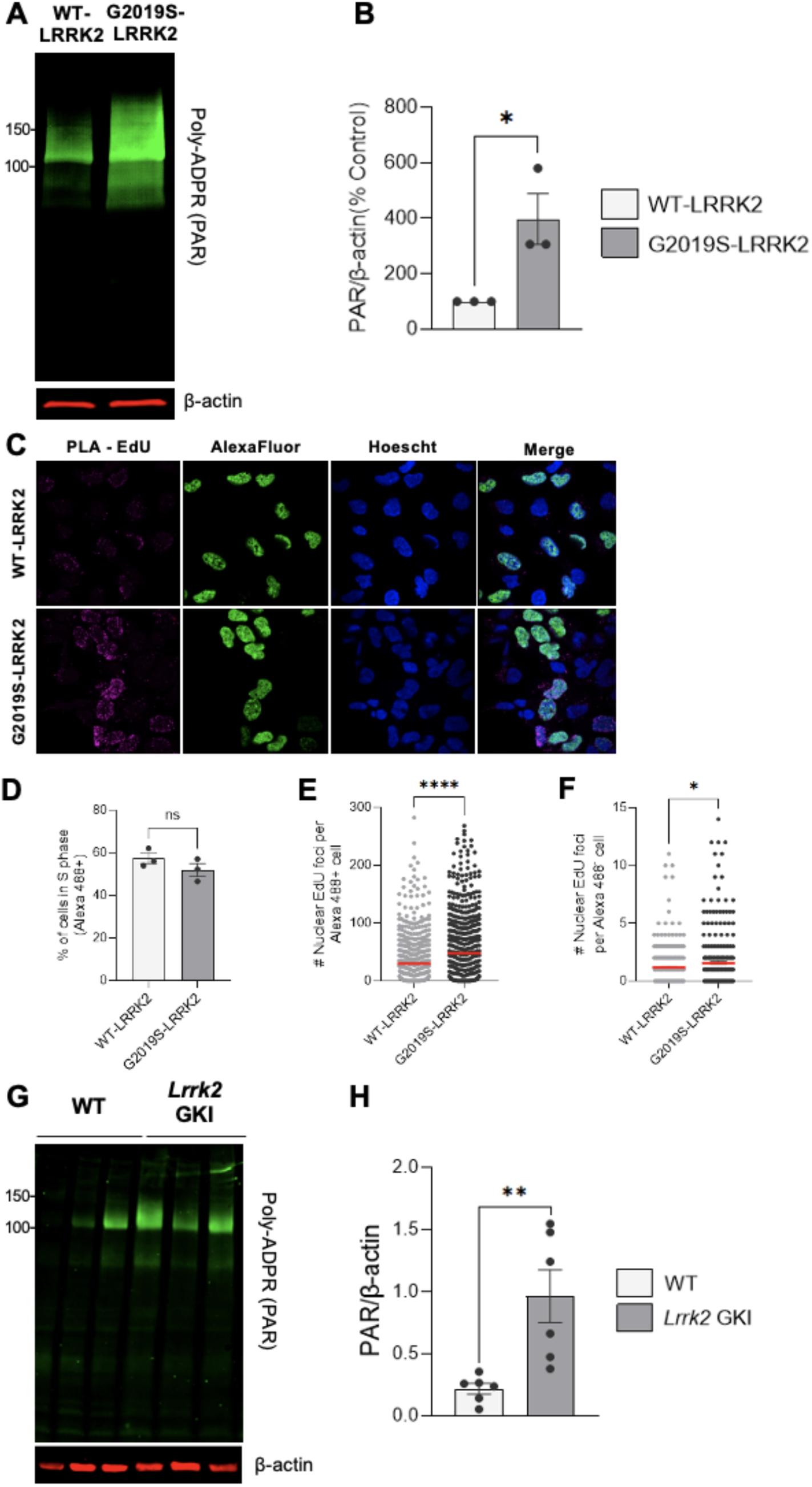
LRRK2 G2019S elevates PAR in cells and mouse midbrain. (**A**) Representative western blot and quantification of WT-LRRK2 and G2019S-LRRK2 expressing cells and assessed for PAR levels and β-actin as a loading control. (**B**) Quantification of PAR reveals higher levels in G2019S-LRRK2 cells relative to WT-LRRK2. (n=3; *p < 0.05 determined by unpaired t-test). (**C**) Representative 60x confocal images of WT-LRRK2 and G2019S-LRRK2 treated with EdU labeled by PLA (magenta) and AlexaFluor 488 (green) and incubated with Hoechst (blue). (**D**) Quantification of percentage of WT-LRRK2 and G2019S-LRRK2 cells in S phase (AlexaFluor 488-positive). Data are from three independent experiments in which approximately 200 total cells were analyzed for each experiment. (n=3, ns, determined by Welch’s t-test) (**E**) Quantification of nuclear PLA-EdU foci per cell in G2019S-LRRK2 relative to WT-LRRK2 cells in S-phase (Alexa Fluor 488-positive). Data are from three independent experiments in which 95-125 cells were analyzed for each experiment. (n=3, ****p < 0.0001, determined by Welch’s t-test) (**F**) Quantification of nuclear PLA-EdU foci per cell in G2019S-LRRK2 relative to WT-LRRK2 cells outside of S-phase (Alexa Fluor 488-negative). Data are from three independent experiments in which 75-100 cells were analyzed for each experiment. (n=3, *p < 0.05, determined by Welch’s t-test) (**G**) Representative western blot of midbrain derived from wild-type (n = 6) and *Lrrk2 GKI* heterozygous mice (n = 6) and PAR levels assessed and β-actin as a loading control. (**H**) Quantification demonstrates increased levels of PAR with *LRRK2* G2019S *in vivo*. (**p < 0.01, determined by unpaired t-test). Data are presented as mean ± SEM.

To assess which PARP is responsible for the G2019S-LRRK2 and WT-LRRK2 induced PAR generation, we treated cells with PARP1, PARP2 or PARP3 selective inhibitors. Treatment with the PARP1-selective inhibitor AZD5305 eliminated PAR signal in both G2019S-LRRK2 and WT-LRRK2 cells (Supplemental Figure 7A, B). Whereas the PARP2- or PARP3- selective inhibitors had no impact on levels of PAR in the G2019S-LRRK2 or WT-LRRK2 cells (Supplemental Figure 7C-F), consistent with our findings in cells expressing endogenous wild-type or mutant LRRK2 (Figure 2C, D).

To evaluate PARP1 signaling *in vivo*, we measured PAR levels in the ventral midbrain of wild-type control and *Lrrk2* G2019S knock-in (GKI) heterozygous mice (Figure 3G) (61). We found that ventral midbrains derived from *Lrrk2* G2019S mice demonstrated increased PAR levels compared with wild-type mice (Figure 3G, H). Collectively, LRRK2 G2019S causes accumulation of endogenous oxidative DNA lesions and robust, PARP1-dependent PAR formation in cells and mouse ventral midbrain.

### LRRK2 G2019S promotes PARP1 hyperactive signaling and selective sensitivity to PARP1-trapping inhibition

Since LRRK2 G2019S caused increased PAR, we next wanted to distinguish whether these elevated levels are due to PARP1 activation or abundance. We first measured *PARP1* transcripts by quantitative RT-PCR and found decreased *PARP1* mRNA levels in LRRK2^G2019S/G2019S^ KI cells relative to wild-type (Figure 4A). Despite reduced *PARP1* RNA expression, PARP1 levels were comparable between LRRK2^G2019S/G2019S^ KI and wild-type cells by immunocytochemistry, with PARP1 localized to the nucleus in both cell lines (Figure 4B, C). Similarly, quantitative western blotting showed equivalent PARP1 protein expression levels in LRRK2^G2019S/G2019S^ KI and wild-type cells (Figure 4D, E). Thus, despite comparable PARP1 protein expression, LRRK2 G2019S cells display markedly elevated PAR, indicating that the PD-linked mutant primarily enhances PARP1 enzymatic activity rather than via abundance, underscoring hyperactivation of PARP1 signaling.

**Figure 4.**
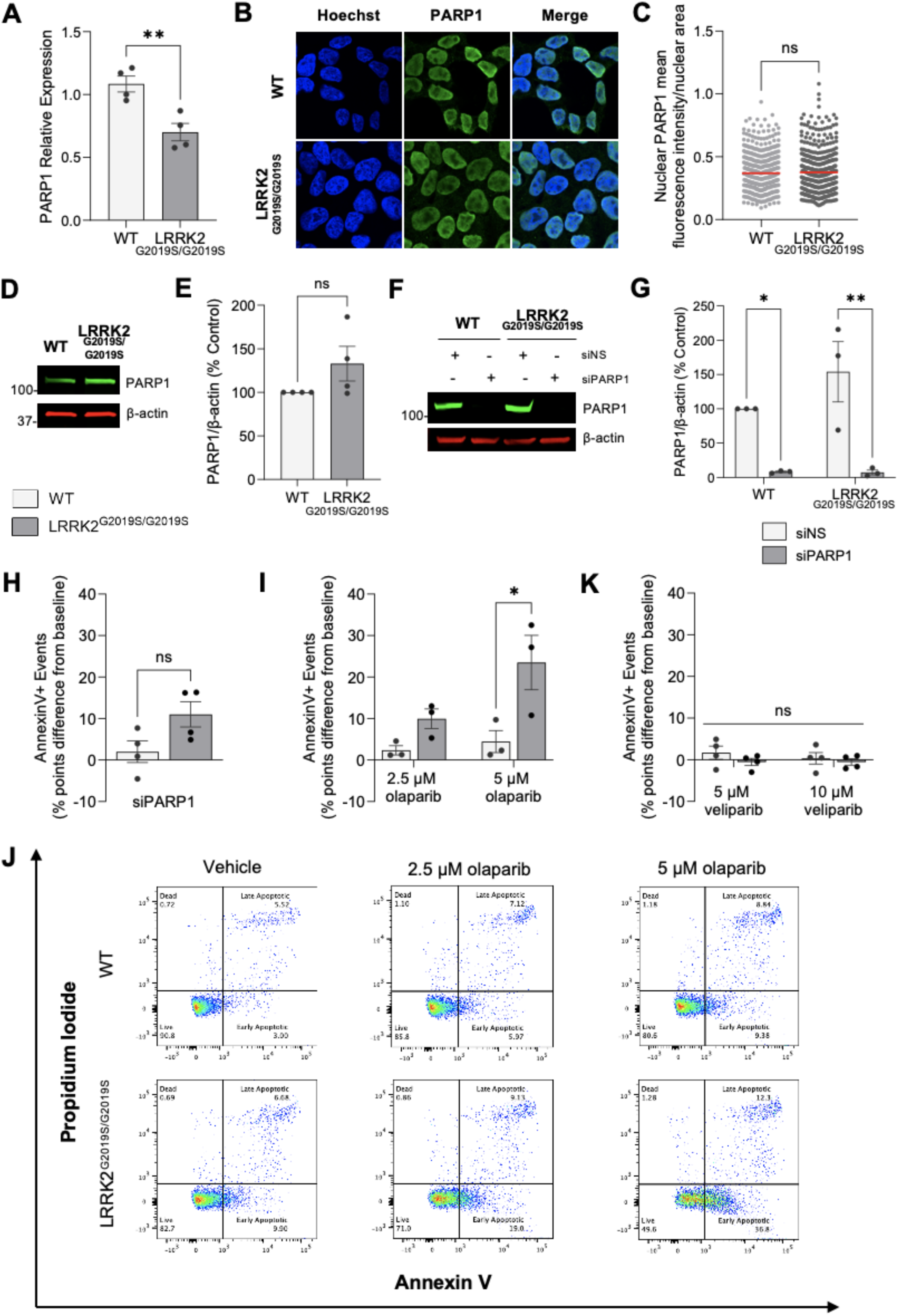
The potent PARP-trapping inhibitor olaparib selectively induces apoptosis in LRRK2^G2019S/S2019S^cells while the weak/non-trapping veliparib does not. (**A**) RNA was collected for qRT-PCR analysis of *PARP1* mRNA expression in wild-type and LRRK2^G2019S/G2019S^ KI cells. Internal control for normalization was GAPDH. (n=4 biological replicates with three technical replicates each, **p < 0.01, determined by unpaired t-test). (**B**) Representative 60X confocal immunofluorescence images of wild-type (WT) and LRRK2^G2019S/G2019S^ KI cells incubated with Hoechst (blue) and PARP1 (green). (**C**) Quantification of nuclear PARP1 signal shows no differences between cell lines. The red bar indicates the mean and each point represents an individual cell. Data are from four independent experiments in which 90–150 nuclei were counted for each experiment. (n=4, ns = non-significant, determined by unpaired t-test). (**D**) Representative western blot of wild-type and LRRK2^G2019S/G2019S^ KI cells assessed for PARP1 levels and β-actin as a loading control. (**E**) Quantification showing similar PARP1 levels in LRRK2^G2019S/G2019S^ KI cells compared to wild-type cells. (n=4, ns=non-significant, determined by unpaired t-test). (**F**) Representative western blot of wild-type and LRRK2^G2019S/G2019S^ KI cells following transient knock-down of siRNA Non-Silencing (siNS) or siRNA against PARP1 (siPARP1) and assessed for PARP1 and β-actin as a loading control. (**G**) Quantification of levels of PARP1 demonstrated that siNS has no impact in either cell line and the siPARP1 was almost fully effective at ablating PARP1 protein. (n=3, **p < 0.01, determined by one-way ANOVA). (**H**) Flow cytometry analysis of annexin V-positive events in wild-type and LRRK2^G2019S/G2019S^ KI cells after transient knock-down of PARP1 compared to scramble siRNA (n=4, ns, determined by unpaired t-test). (**I**) Flow cytometry analysis of annexin V-positive events in wild-type and LRRK2^G2019S/G2019S^ KI cells treated with vehicle or increasing concentrations of olaparib (n=3; *p < 0.05, determined by two-way ANOVA with Bonferroni’s multiple comparison). (**J**) Representative flow cytometry plots of live, apoptotic, and necrotic cell populations in wild-type and LRRK2^G2019S/G2019S^ KI cells treated with vehicle or olaparib. (**K**) Flow cytometry analysis of Annexin V-positive events revealed no differences between wild-type and LRRK2^G2019S/G2019S^ KI cells treated with vehicle or veliparib (n = 4, ns, determined by two-way ANOVA with Bonferroni’s multiple comparison). Data are presented as mean ± SEM.

We next tested the role of PARP1 in LRRK2^G2019S/G2019S^ KI cell viability by targeting PARP1 function via genetic knockdown or pharmacological inhibition. For *PARP1* knockdown, LRRK2^G2019S/G2019S^ KI and wild-type control cells were transiently transfected with either *PARP1*-targeting siRNA (siPARP1) or a non-targeting scramble control (siNS) (Figure 4F, G). PARP1 protein levels were almost completely abolished in both genotypes following *PARP1* siRNA treatment, whereas the scramble siRNA had no effect on baseline PARP1 expression (Figure 4F, G). There were no differences in viability of either LRRK2^G2019S/G2019S^ KI or wild-type cells after transient *PARP1* depletion compared to the scramble siRNA control, as measured by Annexin V/PI staining and flow cytometry (Figure 4H, Supplemental Figure 8A). Transfection with the scramble siRNA control did not impact viability between LRRK2^G2019S/G2019S^ KI and wild-type cells (Supplemental Figure 8B). The lack of a viability effect with genetic depletion of *PARP1* may reflect compensatory activity by PARP2 in PARP1’s absence, which exhibits partially overlapping functions with PARP1 (66,67).

To further assess how loss of PARP signaling affects cell viability, we treated wild-type and LRRK2^G2019S/G2019S^ KI cells with two distinct PARP1/2 inhibitors, olaparib or veliparib, and measured viability by AnnexinV/PI staining. Olaparib exposure induced increased apoptotic cell death selectively in LRRK2^G2019S/G2019S^ KI cells, with no comparable effect in wild-type cells (Figure 4I, J); no PARP1 cleavage fragment was detected under these conditions (Supplemental Figure 9) (68). Since olaparib both inhibits PARP1 catalytic activity and significantly traps PARP1 on DNA (69), we next tested veliparib, a PARP1/2 inhibitor with limited trapping activity. In contrast to olaparib, veliparib had no significant effect on cell viability in either wild-type control or LRRK2^G2019S/G2019S^ KI cells (Figure 4K, Supplemental Figure 8C). Together, these results demonstrate LRRK2 G2019S causes PARP1 signaling hyperactivation and pharmacologic PARP1/2 inhibition causes lethality of LRRK2 G2019S cells in a PARP-trapping–dependent manner.

### Enrichment of chromatin-bound PARP1 and BER factors with PD-linked mutant

Since LRRK2 G2019S cells were exquisitely sensitive to PARP-trapping inhibitors relative to wild-type (Figure 4H-K), we wondered whether PARP1 was differentially associated with chromatin. Nuclear soluble and chromatin-bound fractions were isolated from LRRK2^G2019S/G2019S^ KI and wildtype cells (Figure 5A). Histone H3 (H3) levels were similar within each chromatin enrichment biological replicate between cell lines (Supplementary Figure 10). Nuclear soluble PARP1 levels were identical between wild-type and LRRK2^G2019S/G2019S^ KI cells (Figure 5A, B). However, even in the absence of exogenous stress or PARP1-trapping inhibitors, LRRK2^G2019S/G2019S^ KI cells displayed a significant enrichment of chromatin-bound PARP1 compared with wild-type cells (Figure 5A, C).

**Figure 5.**
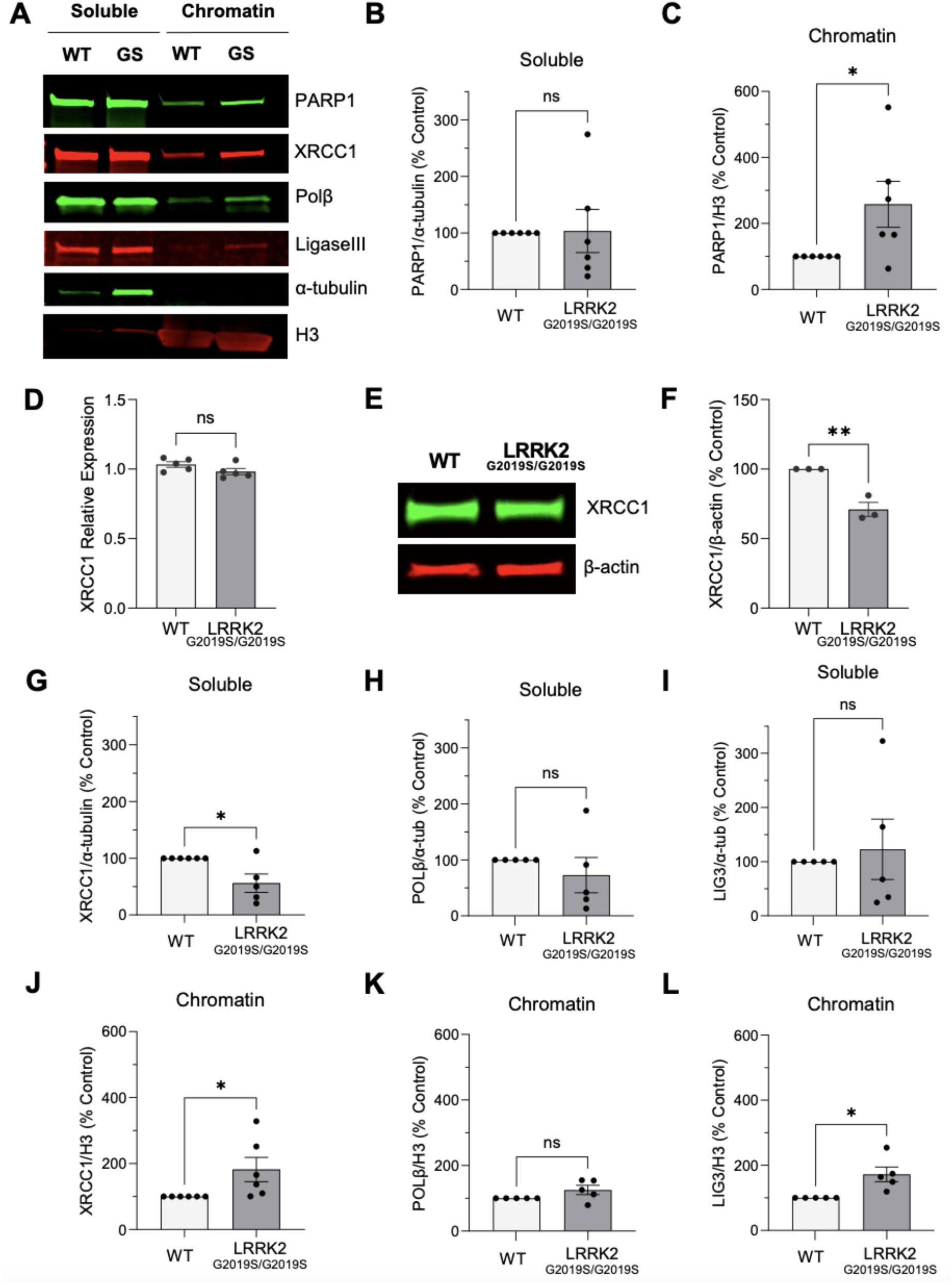
Enrichment of chromatin-bound PARP1 and BER factors in LRRK2^G2019S/S2019S^ cells. (**A**) Representative western blot of soluble and chromatin-enriched fractions in wild-type (WT) and LRRK2^G2019S/G2019S^ KI (GS) cells assessed for PARP1, XRCC1, Pol β, and Ligase III and α-tubulin and H3 as loading controls for soluble and chromatin fractions, respectively. (**B**) Quantification of soluble nuclear PARP1 (n = 6, ns = non-significant, determined by unpaired t-test), and (**C**) chromatin-bound PARP1 levels. (n=6, *p < 0.05, determined by unpaired t-test). (**D**) RNA was collected for qRT-PCR analysis of *XRCC1* mRNA expression in wild-type and LRRK2^G2019S/G2019S^ KI cells. Internal control for normalization was GAPDH. (n=5 biological replicates with three technical replicates each, ns, determined by unpaired t-test). (**E**) Representative western blot of wild-type and LRRK2^G2019S/G2019S^ KI cells assessed for XRCC1 levels and β-actin as a loading control. (**F**) Quantification showing reduced XRCC1 levels in LRRK2^G2019S/G2019S^ KI cells compared to wild-type. (n=6, **p < 0.01, determined by unpaired t-test). (**G**) Quantification of reduced soluble nuclear XRCC1 (n=6, ns or *p < 0.05, determined by unpaired t-test), (**H**) Pol β (n=5, ns), and (**I**) Ligase III levels in wild-type and LRRK2^G2019S/G2019S^ KI cells (n =5, ns or *p < 0.05, determined by unpaired t-test). (**J**) Quantification of enriched chromatin-bound XRCC1, (**K**) unchanged Pol β, or (**L**) increased chromatin associated Ligase III levels in wild-type compared to LRRK2^G2019S/G2019S^ KI cells. (n=3, ns or *p < 0.05, determined by unpaired t-test). Data are presented as mean ± SEM.

Given the increase in chromatin-bound PARP1, we next asked whether other BER scaffolds/enzymes similarly redistribute to chromatin. XRCC1 is a PARP1-dependent scaffold in BER that is recruited to damaged DNA and coordinates downstream BER activities with DNA polymerase β and DNA ligase III. We first measured *XRCC1* transcripts by quantitative RT-PCR and found equivalent *XRCC1* mRNA levels in LRRK2^G2019S/G2019S^ KI cells relative to wild-type (Figure 5D). In contrast, quantitative western blotting showed a modest decrease in XRCC1 protein expression levels in LRRK2^G2019S/G2019S^ KI compared to wild-type cells (Figure 5E, F). While soluble nuclear levels of XRCC1 were decreased (Figure 5G), XRCC1 was enriched in the chromatin-bound fraction derived from LRRK2^G2019S/G2019S^ KI cells (Figure 5A, J).

We further investigated levels of DNA polymerase ꞵ and DNA Ligase III in nuclear-soluble and chromatin fractions derived from wild-type and LRRK2^G2019S/G2019S^ KI cells (Figure 5A). Nuclear-soluble and chromatin-bound DNA polymerase ꞵ was unchanged between genotypes (Figure 5H, K). Finally, there were similar soluble levels of DNA ligase III in LRRK2^G2019S/G2019S^ KI and wild-type cells, but chromatin-bound DNA ligase III was elevated with LRRK2 G2019S (Figure 5I, L). Taken together, these findings indicate active engagement of a subset of BER factors at DNA lesions rather than diffuse or cytosolic signaling.

### SOD/catalase mimetic rescues LRRK2 G2019S dependent PAR accumulation

We next tested whether LRRK2 G2019S-induced PARP1 activation is driven by an ROS-dependent mechanism. EUK-134 is a synthetic superoxide dismutase (SOD) and catalase mimetic (70,71). Treatment with EUK-134 (50 µM, 48 h) reduced PAR levels in LRRK2^G2019S/G2019S^ KI cells compared to those of wild-type controls (Figure 6A, B). EUK-134 exposure had no impact on wild-type PAR basal levels (Figure 6A, B). To further define which antioxidant activity is required to normalize PARP1 activation, we used EUK-8, a related SOD/catalase mimetic with substantially weaker catalase activity (71). EUK-8, even at concentrations up to 50 µM, did not significantly affect PAR levels in either LRRK2^G2019S/G2019S^ KI or wild-type cells, suggesting that the combination of the SOD and catalase activity is required to rescue the PAR phenotype (Figure 6C, D).

**Figure 6.**
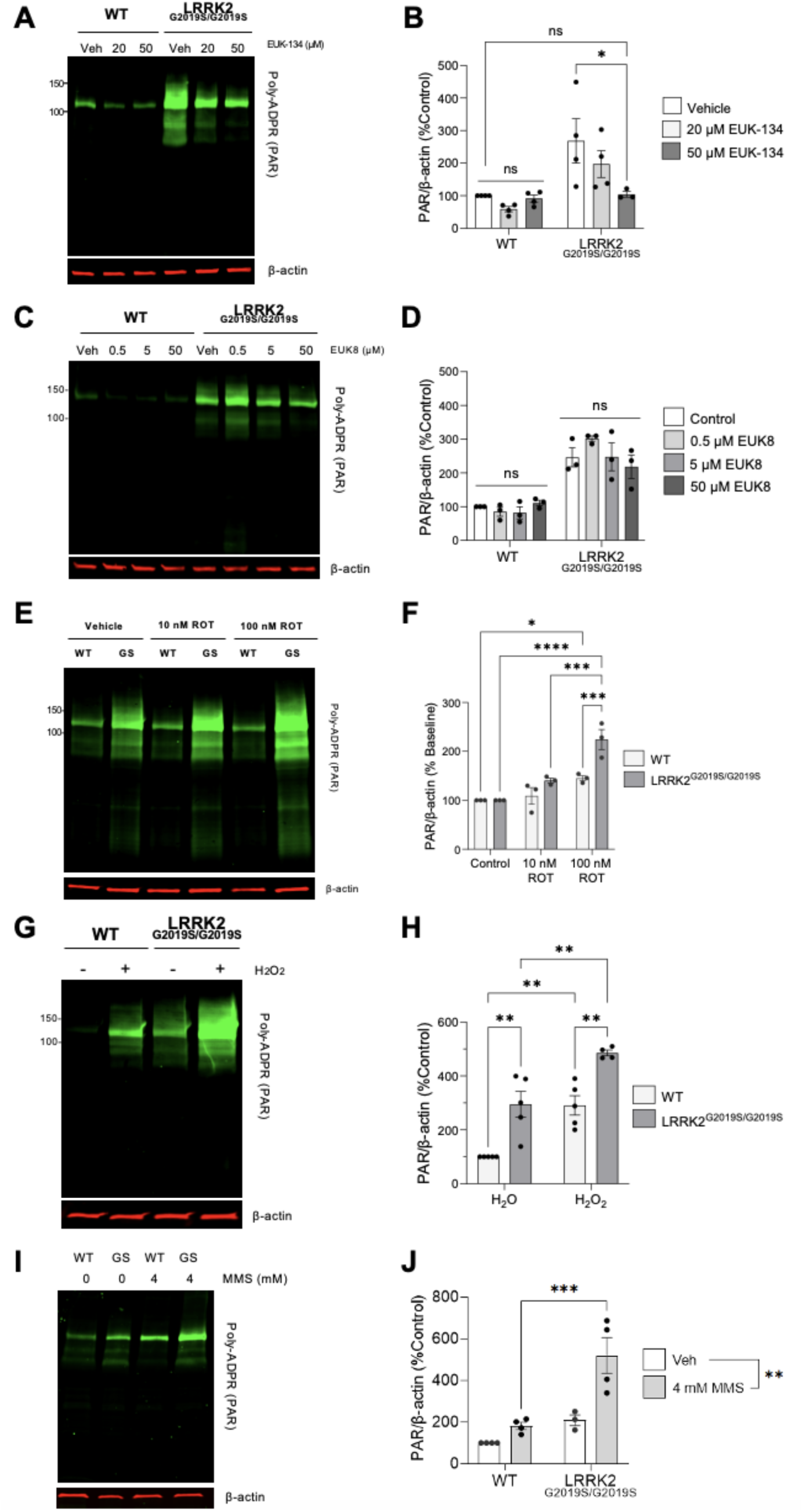
EUK-134 rescues PARP1 activation in LRRK2^G2019S/G2019S^ cells. (**A**) Representative western blot of wild-type and LRRK2^G2019S/G2019S^ KI cells treated with vehicle or EUK-134 for 48 h and assessed for PAR levels and β-actin as a loading control. (**B**) Quantification showing treatment with 50uM of EUK-134 rescued LRRK2 G2019S-induced PAR levels to wild-type baseline. (n=4, *p < 0.05, determined by two-way ANOVA with Bonferroni’s multiple comparison). (**C**) Representative western blot of wild-type and LRRK2^G2019S/G2019S^ KI cells treated with vehicle or EUK-8 for 48 h and assessed for PAR levels and β-actin as a loading control. (**D**) Quantification showing treatment with EUK-8 has no impact on PAR levels in either cell line. (n=3, ns, determined by two-way ANOVA with Bonferroni’s multiple comparison). (**E**) Representative western blot of wild-type and LRRK2^G2019S/G2019S^ KI cells treated with vehicle or rotenone and assessed for PAR levels and β-actin as a loading control. (**F**) Quantification showing rotenone treatment increased PAR levels particularly at higher concentrations in LRRK2^G2019S/G2019S^ KI cells (n=3, *p < 0.05, ***p < 0.001, ****p < 0.0001, determined by two-way ANOVA with Bonferroni’s multiple comparison). (**G**) Representative western blot of wild-type and LRRK2^G2019S/G2019S^ KI cells treated with vehicle or acute H_2_O_2_ and assessed for PAR levels and β-actin as a loading control. (**H**) Quantification showing H_2_O_2_ treatment increased PAR levels in both genotypes (n=5, **p < 0.01, determined by two-way ANOVA with Bonferroni’s multiple comparison). (**I**) Representative western blot of wild-type and LRRK2^G2019S/G2019S^ KI cells treated with vehicle or MMS and assessed for PAR levels and β-actin as a loading control. (**J**) Quantification showing MMS treatment increased PAR levels in LRRK2^G2019S/G2019S^ KI cells (n=4, **p < 0.01, ***p < 0.001, determined by two-way ANOVA with Bonferroni’s multiple comparison). Data are presented as mean ± SEM.

We next examined whether increasing mitochondrial ROS further stimulates PARP1 activation and if LRRK2 G2019S cells are selectively vulnerable to this stress. Rotenone inhibits complex I of the electron transport chain (ETC), and is a PD-relevant toxicant that increases mitochondrial ROS, mtDNA damage and recapitulates many of the core features of PD in cell and animal models (72–75). Rotenone caused an increase in PAR accumulation in both wild-type and LRRK2^G2019S/G2019S^ KI cells, but LRRK2 G2019S cells demonstrated enhanced PAR generation with rotenone treatment. In LRRK2^G2019S/G2019S^ KI cells, rotenone exposure produced an additional ∼two-fold increase in PAR levels over the already elevated baseline (Figure 6E, F), whereas wild-type cells challenged with rotenone treatment exhibited a ∼45% increase in PAR (Figure 6E, F). These findings indicate that mitochondrial ROS synergistically amplifies LRRK2 G2019S-associated PARP1 activation.

Given that PD-relevant mitochondrial insults that generate ROS exacerbate the production of PAR, we investigated whether exogenous DNA-damaging agents that generate oxidative and alkylation lesions processed by BER, magnifies PARP1 activation. Incubation with hydrogen peroxide (400 µM, 10 min) caused an increase in PAR generation in LRRK2^G2019S/G2019S^ KI cells compared to wild-type (Figure 6G, H). Exposure to the alkylative agent MMS (4 mM, 25 min) similarly magnified PARP1 activation in LRRK2^G2019S/G2019S^ KI cells (Figure 6I, J). Similar findings of elevated PAR levels associated with G2019S-LRRK2 cells following MMS exposure were quantified by flow cytometry (Supplementary Figure 11). Although gene expression is tightly regulated, abortive or stalled events during transcription can cause genome instability and enhancers are hotspots for endogenous single-strand breaks in neurons (45). We observed that LRRK2-dependent PARP1 activation is transcription independent, as inhibition of RNA Polymerase II with 5,6-Dichloro-1-β-D-ribofuranosylbenzimidazole (DRB) had no effect on PAR levels (Supplemental Figure 12). These data support a heightened PARP1 response to and/or repair imbalance of BER-relevant base damage associated with the PD-linked LRRK2 G2019S mutation.

## DISCUSSION

To our knowledge, this study provides the first direct evidence that the PD-linked LRRK2 G2019S mutation causes endogenous oxidative damage to the nuclear genome. We detected increased oxidative base lesions and elevated PAR levels, which indicates PARP activation in response to BER intermediates and DNA-strand breaks, and is consistent with persistent, unresolved endogenous genotoxic stress. These results extend prior reports of LRRK2 G2019S-associated mitochondrial DNA damage, and align with studies linking LRRK2 to the ATM-mediated DNA damage response (DDR) (28,76). Supporting a broader connection between PD-linked proteins and nuclear genome instability, multiple studies implicate α-synuclein in nuclear DNA repair. Specifically, oxidized or mislocalized α-synuclein (including familial A53T mutants) can impair nuclear DNA repair and increase vulnerability to DNA damage (77–84). More generally, DNA damage and DDR markers accumulate in the brains of individuals with PD, suggesting defective genome maintenance contributes to neuronal dysfunction and loss (22,24,59,79,85–87). Since LRRK2 can affect nuclear processes despite limited evidence for stable nuclear localization (76,88–90), future work may define whether LRRK2 effects on genome homeostasis are a result of direct nuclear mechanisms and/or via indirect cytoplasmic/mitochondrial pathways.

Accumulating evidence implicates PARP1 and PAR in PD pathogenesis (57,91,92). We observed PARP1-dependent hyperactivation in multiple LRRK2 G2019S models. Our findings are consistent with studies showing that pathologic α-synuclein activates PARP1, PAR levels are elevated in brains derived from PD patients, and PAR promotes α-synuclein fibrillization and toxicity *in vitro* and *in vivo* (59). Genetic or pharmacologic PARP1 inhibition prevented α-syn PFF-induced neuronal death and propagation in some reports (59), although another study found only modest effects of PAR on α-synuclein fibrillization and no protection against α-synuclein inclusion formation or dopaminergic cell loss *in vivo* (93). Prior studies demonstrated PARP inhibition is cytoprotective in α-synuclein and MPP^+^ *in vitro* models; and PARP1 activation mediates dopaminergic neuron loss in *in vivo* MPTP toxin models of parkinsonism (94,95).

Despite LRRK2 G2019S-induced increased PARP1 signaling, PARP1 inhibition did not rescue viability and instead increased sensitivity to cell death, suggesting that PARP1 initially plays a protective role in this context. While PARP1 recruitment to acute DNA damage promotes repair, chronic activation under sustained oxidative stress can exhaust NAD^+^/ATP and trigger parthanatos (96,97). Although we did not detect mitochondrial PAR, it is possible that exogenous stress is necessary to drive parthanatos in our LRRK2 mutant models. Dopaminergic neurons are particularly vulnerable to oxidative lesions and accumulate single-strand DNA breaks that activate PARP1 (79,98,99), and persistent PARP1 activation can shift from protective repair to a driver of metabolic failure. Interventions that boost NAD^+^ (e.g., nicotinamide riboside) or limit PARP activity have been demonstrated to improve mitochondrial function and survival in PD models and safety in early human clinical trials, and represent potential complementary strategies to mitigate PARP-driven toxicity (59,100,101). Together, these data suggest a duality of PARP1 signaling, where PARP1 initially facilitates repair and survival in LRRK2 G2019S cells but may become deleterious with persistent activation or aging–a mechanism that may be distinct from α-synuclein mediated toxicity. However, longer-term studies are needed to further understand the multifaceted roles for PARP1, in particular in neurons and glia, and to define the balance between protective and pathogenic PARP1 roles, connecting DNA damage, metabolic stress and pathology in PD.

Carriers of PD-linked *LRRK2* mutations are at a higher risk of developing breast cancer (102–104). Our data indicate that LRRK2 G2019S drives persistent oxidative base lesions and PARP1 activation, consistent with potential defects in BER/SSBR and impaired resolution of oxidative intermediates. Such repair insufficiency creates actionable vulnerabilities in cancer. Specifically, tumors with BER/SSBR defects exhibit synthetic-lethal sensitivity to PARP inhibitors (105). Furthermore, these tumors are sensitive to agents that generate persistent SSBs, such as topoisomerase-I poisons (e.g., camptothecins), where the choice of combination therapies is guided by PARP trapping and catalytic inhibition (106). Combined targeting of complementary DDR pathways can convert unrepaired SSBs or trapped PARP-DNA complexes into lethal DSBs (i.e., ATR inhibitors synergize with PARP inhibitors) (107). Similarly, Polθ/POLQ inhibition is synthetically lethal with HR defects and potentiates PARP inhibitor responses (108), and ATM functions upstream in the response to PARP inhibitor-induced collapsed forks (109). Consistent with a PARP-targetable liability, LRRK2 G2019S cells are more vulnerable to the PARP1 inhibitor olaparib compared to veliparib, which is likely due to differences in PARP-trapping potency (110). PARP1 hyperactivation can be a feature of both defective BER and HR (111). Increased PAR accumulation has been reported when BER is impaired (112,113). Furthermore, BER dysfunction and loss of XRCC1 lead to excessive PARP1 engagement/trapping and increased sensitivity to PARP inhibition (114–117). Future experiments measuring BER and HR efficiency and fidelity are therefore needed to determine the underlying LRRK2-mediated repair defect(s) to rationalize PARP-based or combination DDR therapeutic strategies.

## SUPPLEMENTARY DATA

Supplementary Data are available online.

## ACKNOWLEDGEMENTS

We thank all members of the Sanders laboratory for technical assistance and discussions. We are also grateful to the Duke Light Microscopy Core Facility (LMCF) and Duke Flow Cytometry Shared Resource for resources and assistance. Finally, we would like to thank Dr. Shan Zha for providing PARP1 plasmids.

## AUTHOR CONTRIBUTIONS

J.L. conceptualized the research studies, conducted the cell viability assays, *in vitro* and *in vivo* western blots, fractionations, immunofluorescence, data analyses, data visualization, and drafted, wrote, and revised the manuscript. C.P.G. conceptualized the research studies, conducted western blots for PAR levels in LRRK2 overexpressing cell lines, performed analyses, and reviewed and edited the manuscript. T.R. supervised MIRA assays and analyses and reviewed and edited the manuscript. I.B. supervised qRT-PCR experiments and reviewed and edited the manuscript. C.C. performed MIRA assays and analyses and reviewed and edited the manuscript. C.M. conducted qRT-PCR experiments and reviewed and edited the manuscript. L.M. conducted flow cytometry measuring PAR in LRRK2 overexpressing cell lines and reviewed and edited the manuscript. R.C. performed western blots with PARP inhibitors in LRRK2 overexpressing cell lines and reviewed and edited the manuscript. V.H. performed western blots for PAR levels after exposure to MMS and reviewed and edited the manuscript. N.R.G. conducted the oxRADD assays and analyses and reviewed and edited the manuscript. E.F. reviewed and edited the manuscript. L.H.S. conceptualized the research studies, supervised all experiments and analyses, wrote and revised the manuscript, and acquired funding.

## FUNDING

This work was supported by National Institutes of Health [grant numbers RO1NS119528 to L.H.S., RO1NS119528-03S2 to L.H.S.].

## CONFLICT OF INTEREST

None declared.

## SUPPLEMENTAL FIGURES

**Supplementary Figure 1.**
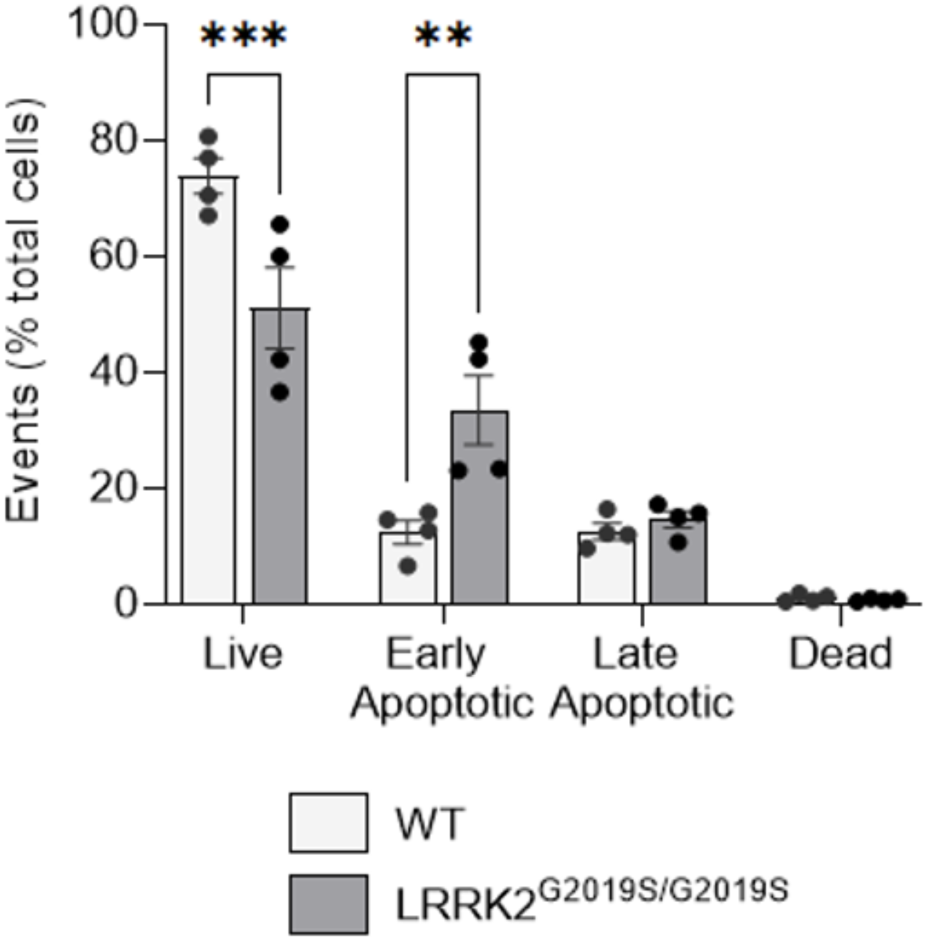
LRRK2^G2019S/G2019S^ cells have decreased viability compared to wild-type. Flow cytometry analysis of live, apoptotic, and necrotic cells in wild-type and LRRK2^G2019S/G2019S^ cells. (n=4, **p < 0.01, ***p < 0.001 determined by two-way ANOVA with Bonferroni’s multiple comparison). Data are presented as mean ± SEM.

**Supplementary Figure 2.**
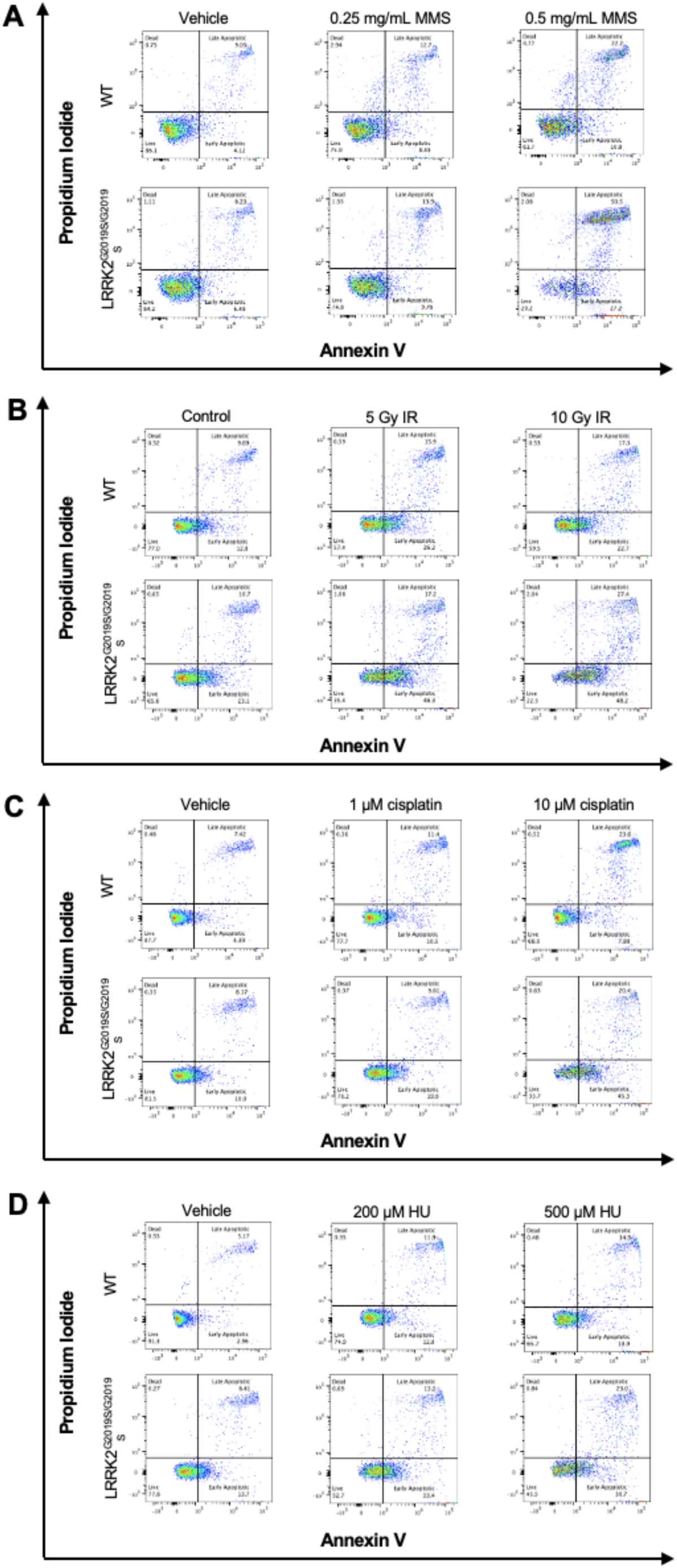
The LRRK2 G2019S mutation sensitizes cells to DNA damaging agents *in vitro*. Representative flow cytometry plots of live, apoptotic, and necrotic cells in wild-type and LRRK2^G2019S/G2019S^ KI cells exposed to (**A**) methyl methanesulfonate (MMS), (**B**) ionizing radiation (IR), (**C**) cisplatin, or (**D**) hydroxyurea (HU) and stained for annexin V/propidium iodide (PI).

**Supplementary Figure 3.**
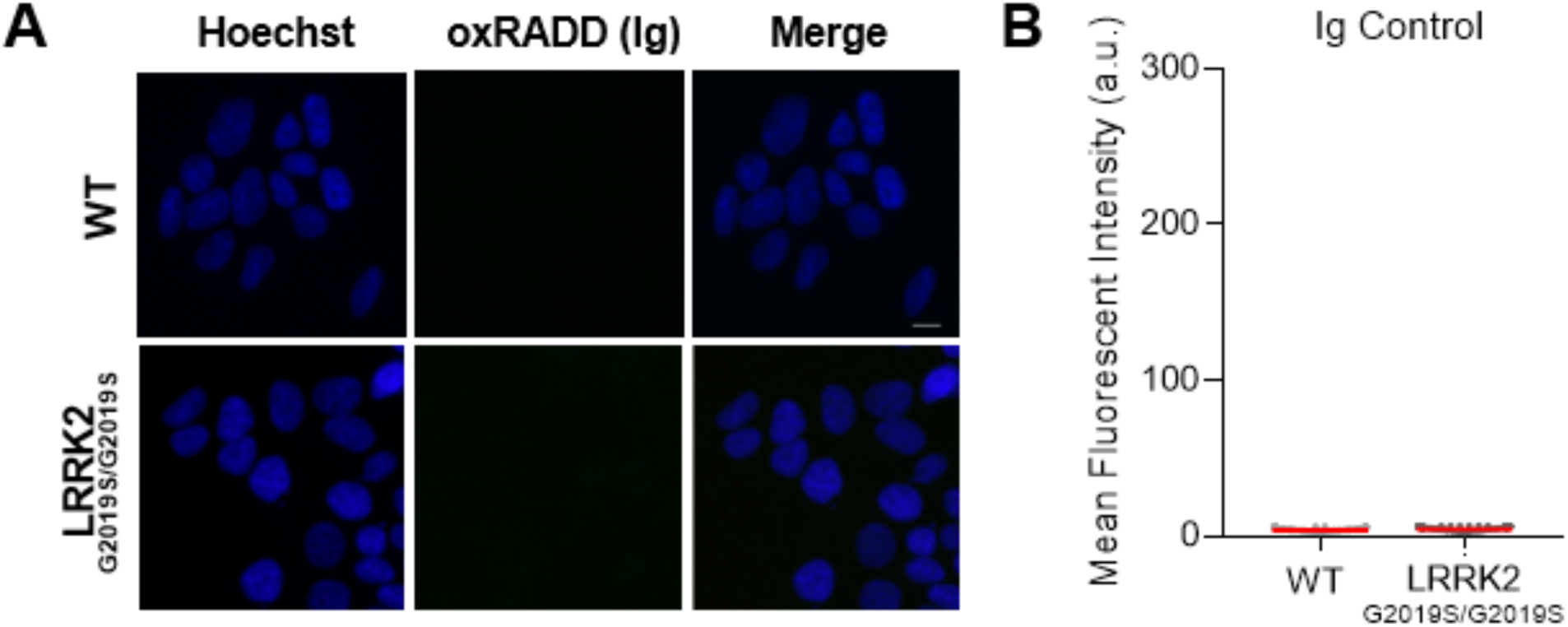
Isotype controls in wild-type and LRRK2^G2019S/G2019S^ cells for the oxRADD assay. (**A**) Representative 20X wide-field images of wild-type (WT) and LRRK2^G2019S/G2019S^ KI cells incubated with Hoechst (blue) and isotype control (green). (**B**) Quantification of IgG control (n=3). Approximately 1700-2000 total cells were analyzed from three independent experiments. Data are presented as mean ± SEM.

**Supplementary Figure 4.**
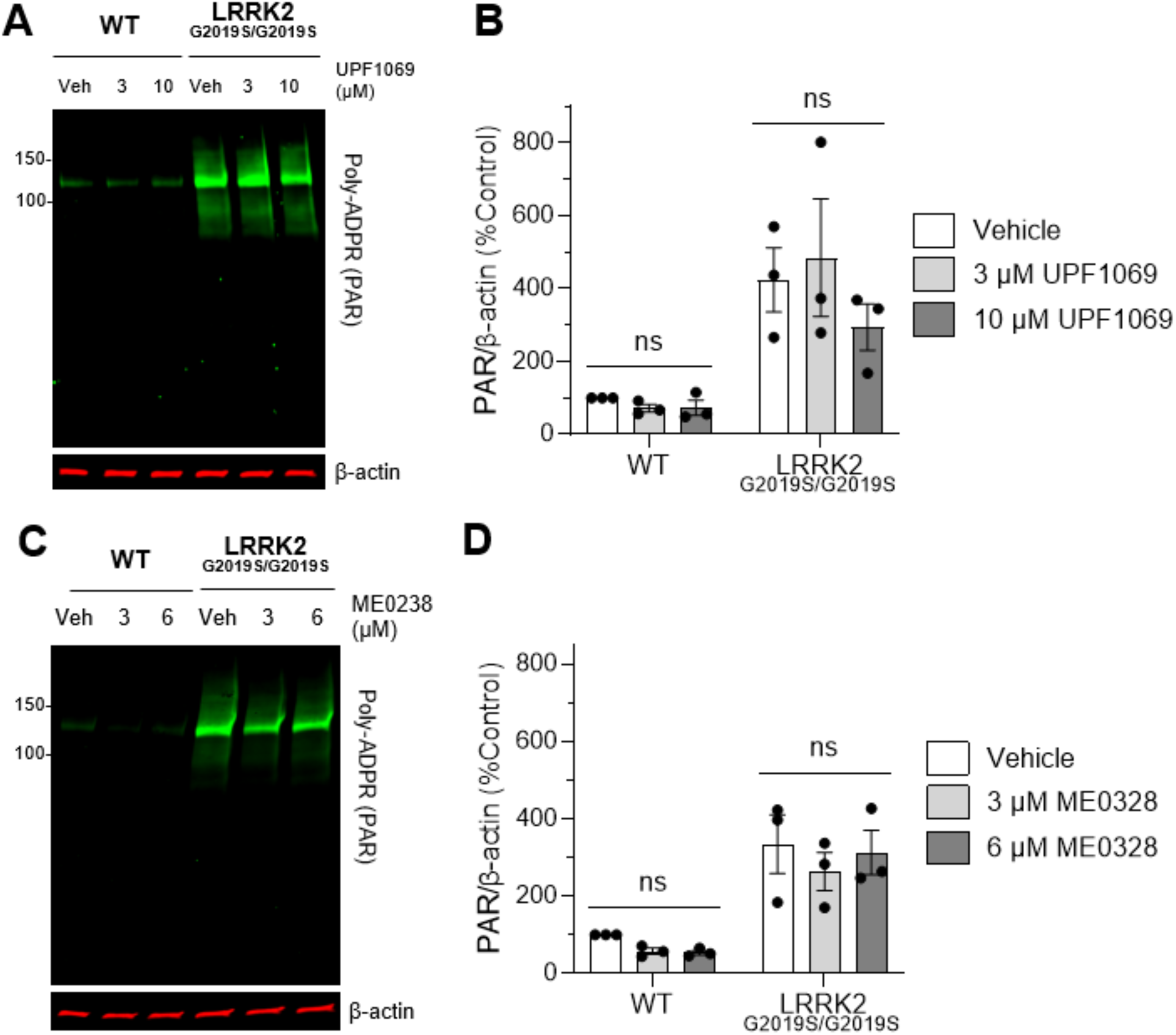
LRRK2 G2019S-mediated PAR accumulation *in vitro* is not PARP2-or PARP3-dependent. (**A**) Representative western blot of wild-type and LRRK2^G2019S/G2019S^ KI cells treated with vehicle or the PARP2-selective inhibitor UPF1069 (3 and 10 µM) and assessed for PAR and β-actin as a loading control. (**B**) No differences in quantification of PAR levels with treatment in either cell line. (n=3, ns, determined by two-way ANOVA with Bonferroni’s multiple comparison). (**C**) Representative western blot of wild-type and LRRK2^G2019S/G2019S^ KI cells treated vehicle or the PARP3-selective inhibitor ME0238 (3 and 6 µM) and assessed for PAR and β-actin as a loading control. (**D**) No differences in quantification of PAR levels with treatment in either cell line. (n=3, ns, determined by two-way ANOVA with Bonferroni’s multiple comparison). Data are presented as mean ± SEM.

**Supplementary Figure 5.**
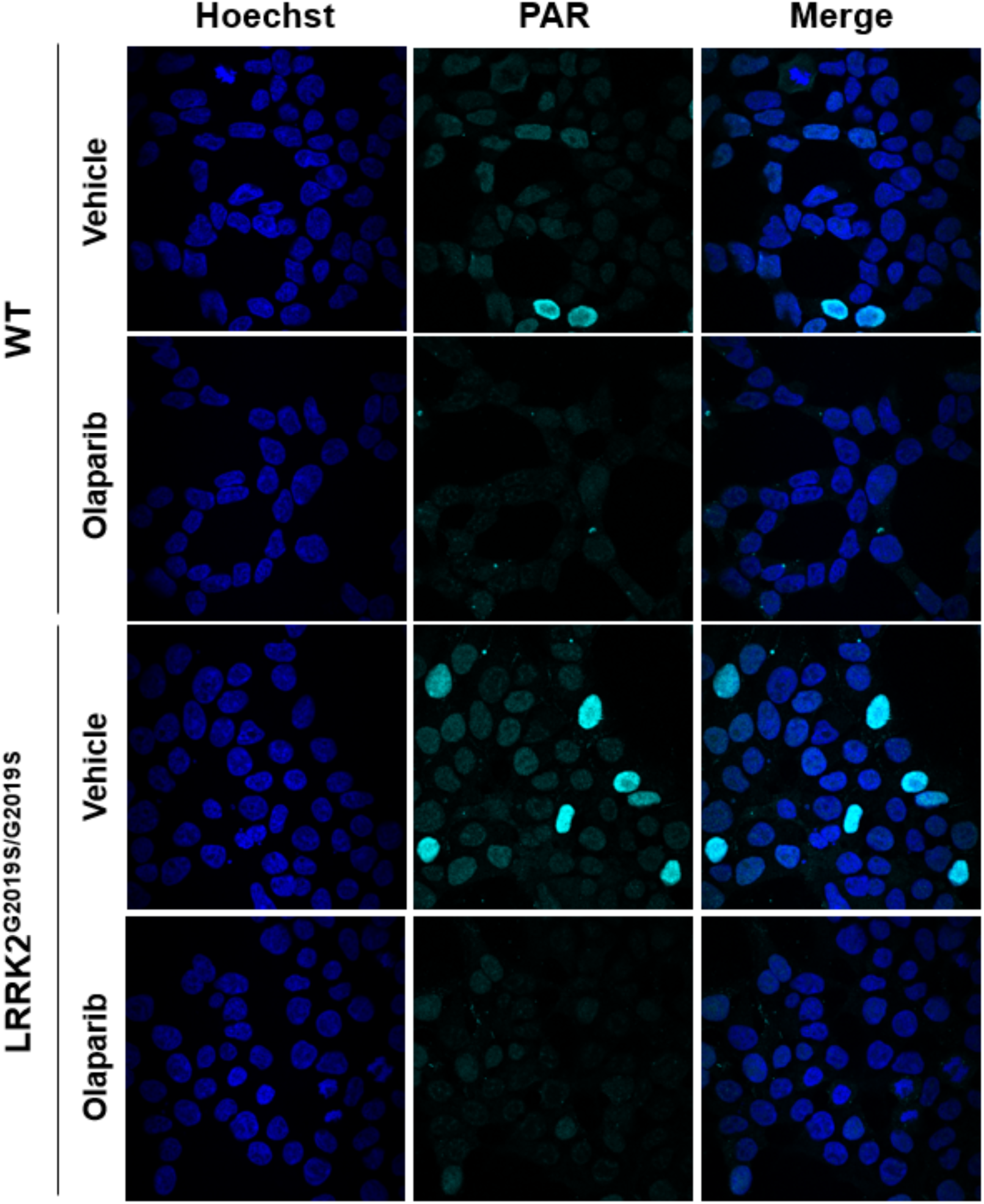
PAR signal is abrogated with olaparib treatment in wild-type and LRRK2^G2019S/G2019S^ cells. Representative 60X confocal fluorescence images of wild-type and LRRK2^G2019S/G2019S^ cells incubated with vehicle or olaparib (10 µM) for 1 h.

**Supplementary Figure 6.**
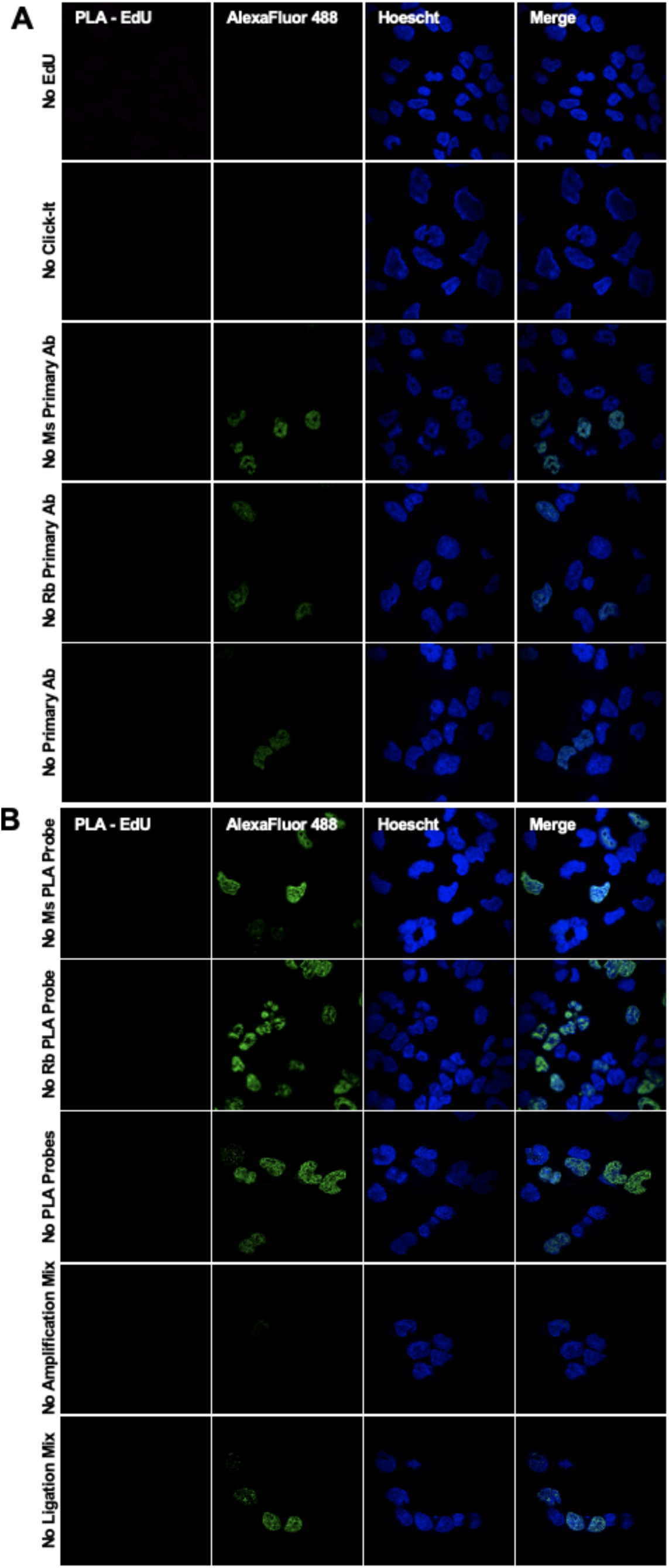
Proximity ligation assay controls. To assess for non-specific or potential background staining in PLA staining, we performed the MIRA protocol in (**A**) WT-LRRK2 cells without the addition of EdU, omitting the steps of the Click-IT reaction, or no addition of the individual mouse or rabbit primary antibody, or primary antibodies, and additional controls that included (**B**) lack of individual mouse or rabbit PLA probes, or both PLA probes, or the PLA amplification or ligation mix was excluded.

**Supplementary Figure 7.**
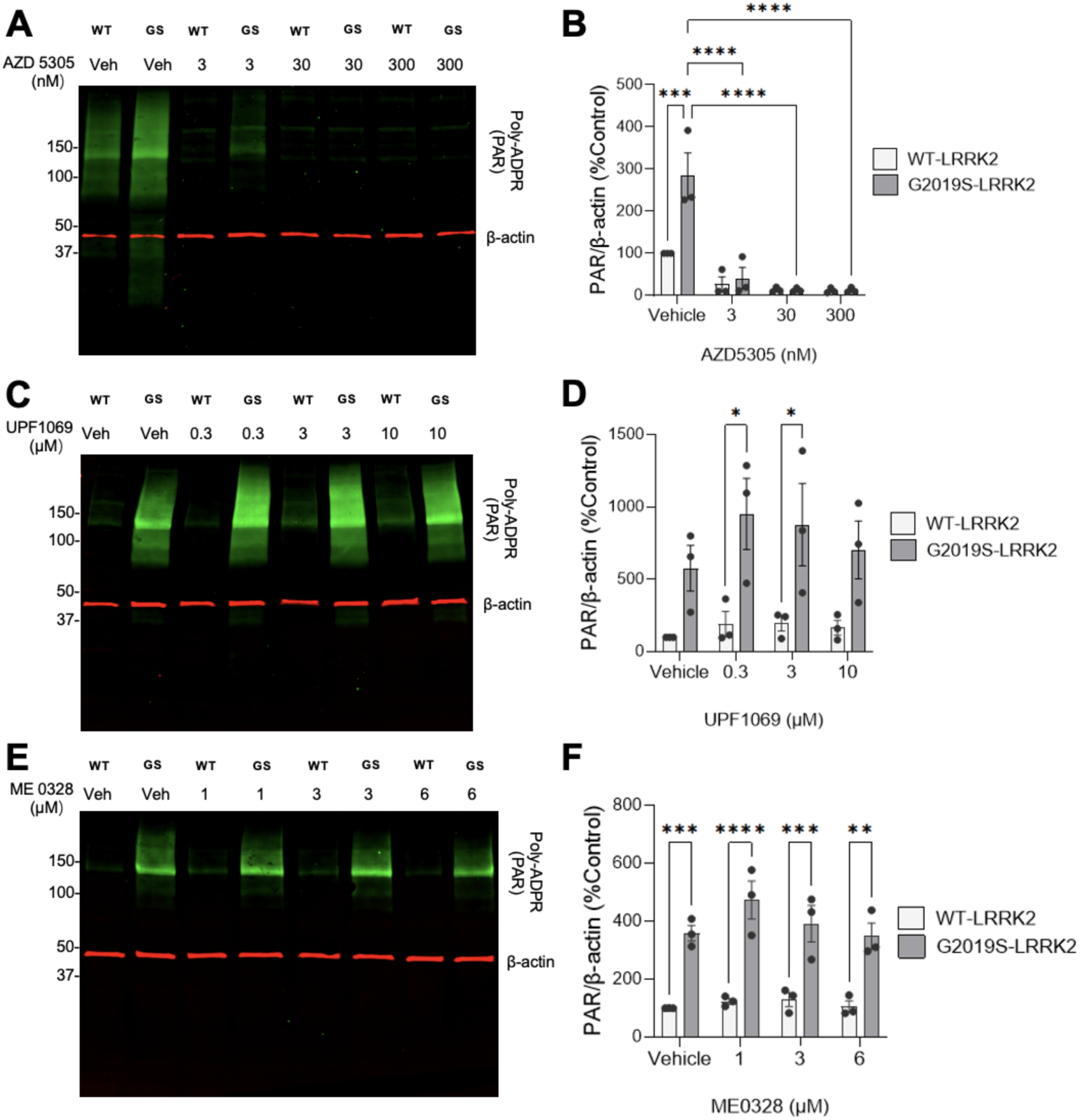
G2019S-LRRK2 mediated PAR accumulation is PARP1-dependent *in vitro*. (**A**) Representative western blot of WT-LRRK2 and G2019S-LRRK2 cells treated with vehicle or the PARP1-selective inhibitor AZD5305 (3, 30 and 300 nM) and assessed for PAR and β-actin as a loading control. (**B**) Quantification demonstrated PAR levels are PARP1-dependent. (n =3, ***p < 0.001, ****p < 0.0001, determined by two-way ANOVA with Bonferroni’s multiple comparison). (**C**) Representative western blot of WT-LRRK2 and G2019S-LRRK2 cells treated with vehicle or the PARP2-selective inhibitor UPF1069 (0.3, 3 and 10 µM) and assessed for PAR and β-actin as a loading control. (**D**) No differences in quantification of PAR levels with treatment in either cell line. (n=3, ns, determined by two-way ANOVA with Bonferroni’s multiple comparison). (**E**) Representative western blot of WT-LRRK2 and G2019S-LRRK2 cells treated with vehicle or the PARP3-selective inhibitor ME0238 (1, 3 and 6 µM) and assessed for PAR and β-actin as a loading control. (**F**) No differences in quantification of PAR levels with treatment in either cell line. (n=3, ns, determined by two-way ANOVA with Bonferroni’s multiple comparison). Data are presented as mean ± SEM.

**Supplementary Figure 8.**
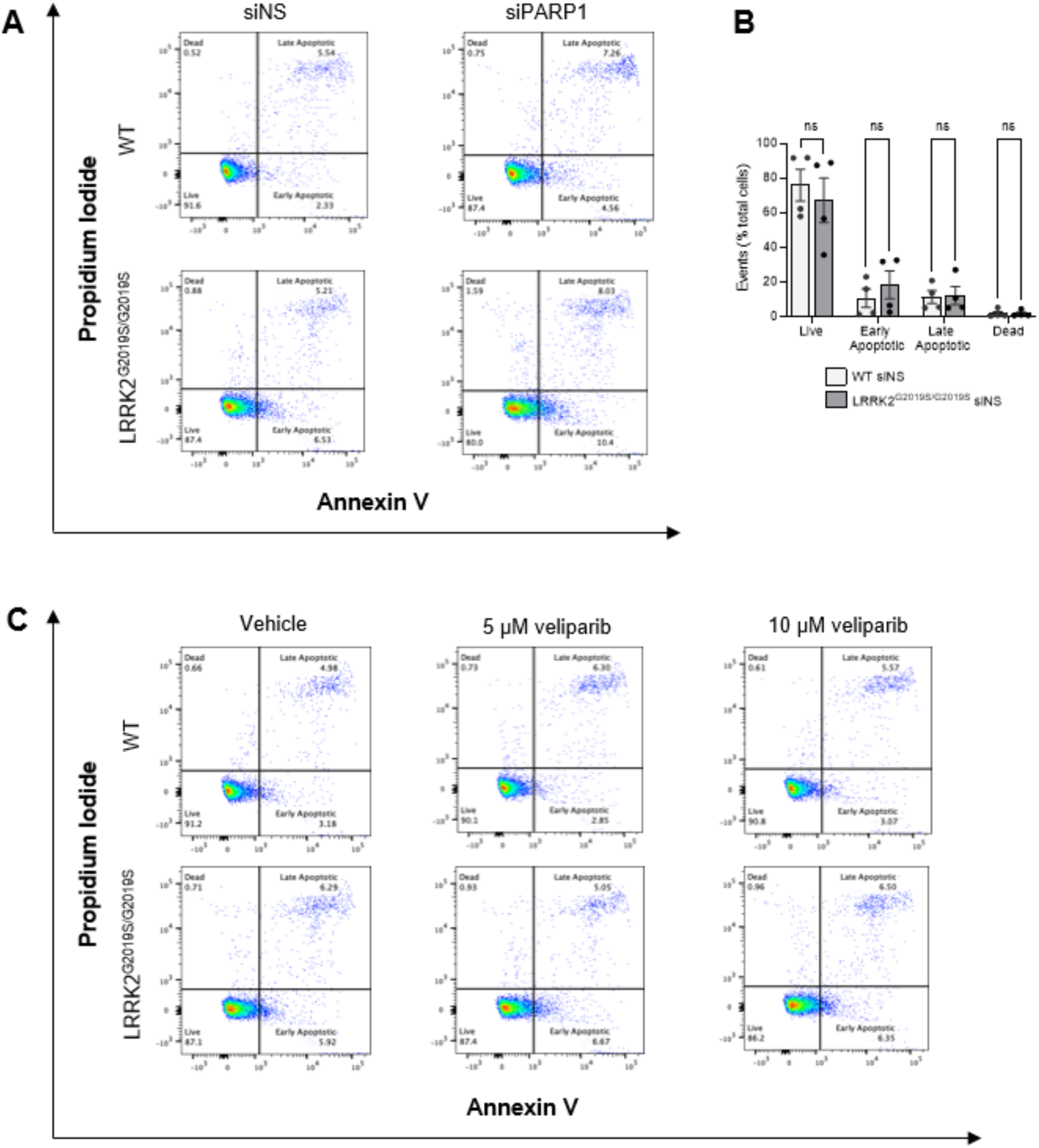
Viability of wild-type and LRRK2^G2019S/S2019S^ cells are similar following either PARP1 knock-down or inhibition with veliparib. (**A**) Representative flow cytometry plots of wild-type and LRRK2^G2019S/G2019S^ KI cells stained with annexin V/propidium iodide (PI) after transient knock-down with siNS or siPARP1 (48 h). (**B**) Live, apoptotic, and necrotic populations of wild-type and LRRK2^G2019S/G2019S^ KI cells after transfection with scramble siRNA (siNS). (n=4, ns = non-significant, determined by two-way ANOVA with Bonferroni’s multiple comparison). (**C**) Representative flow cytometry plots of wild-type and LRRK2^G2019S/G2019S^ KI cells treated with vehicle or veliparib and stained with annexin V/PI. Data are presented as mean ± SEM.

**Supplementary Figure 9.**
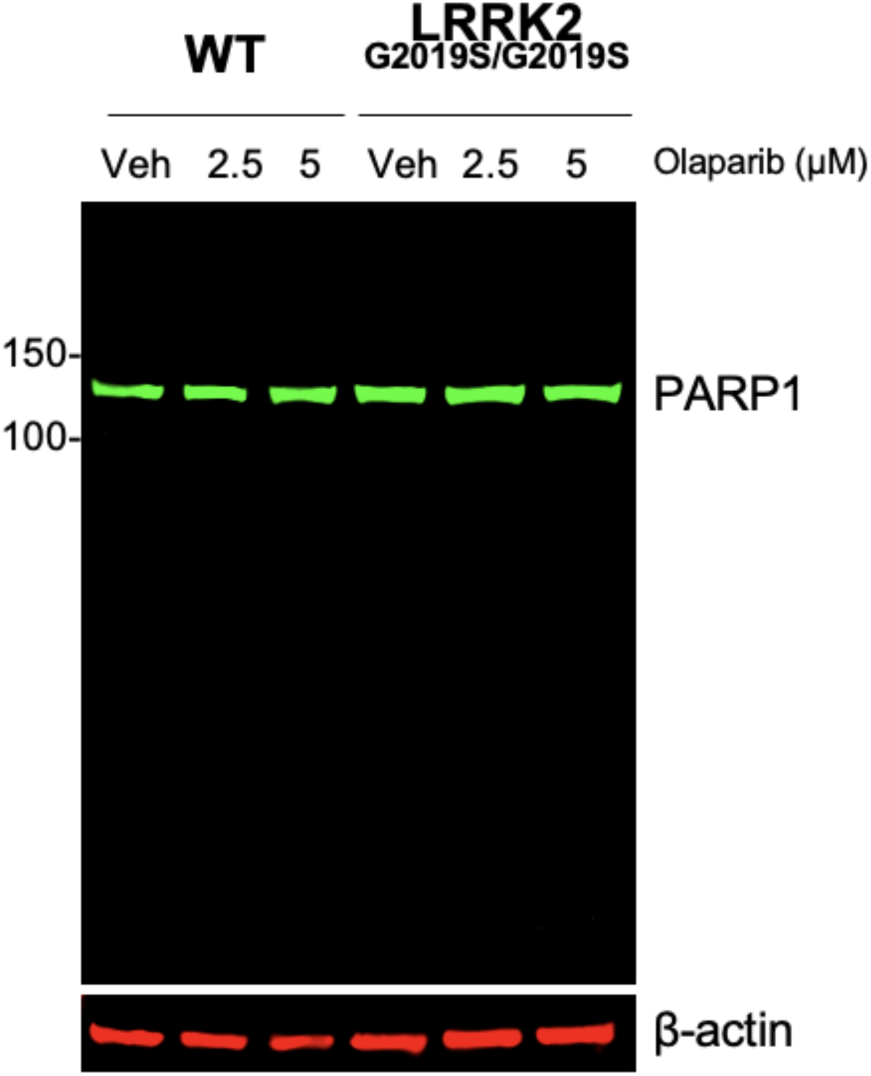
LRRK2^G2019S/G2019S^ KI and wild-type cells do not exhibit PARP1 cleavage after treatment with olaparib. Representative western blot of wild-type (WT) and LRRK2^G2019S/G2019S^ KI cells treated with vehicle or olaparib (2.5 and 5 µM) for 48 h and assessed for PARP1 and β-actin as loading control.

**Supplementary Figure 10.**
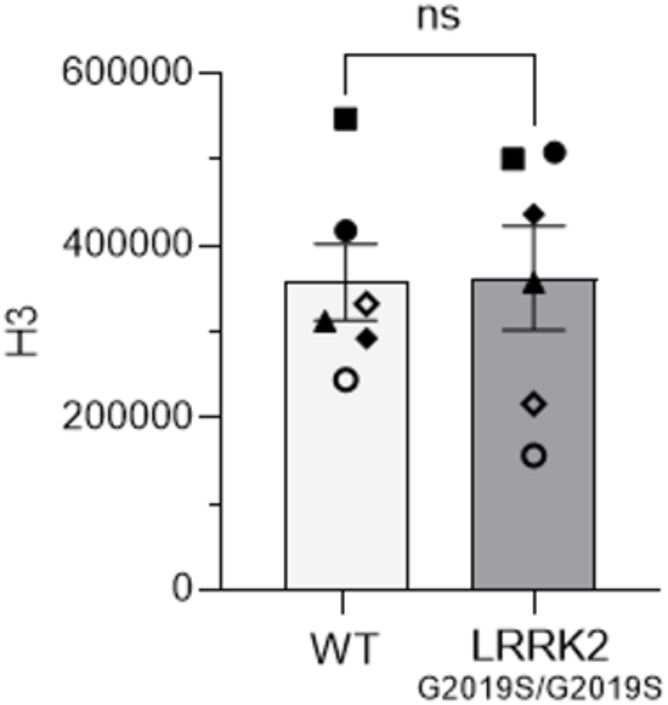
H3 levels are equivalent in chromatin-enriched fractions between LRRK2^G2019S/S2019S^ KI and wild-type cells. Quantification of H3 levels in chromatin-enriched fractions from wild-type (WT) and LRRK2^G2019S/G2019S^ KI cells. (n=6, ns, determined by an unpaired t-test). Data are presented as mean ± SEM.

**Supplementary Figure 11.**
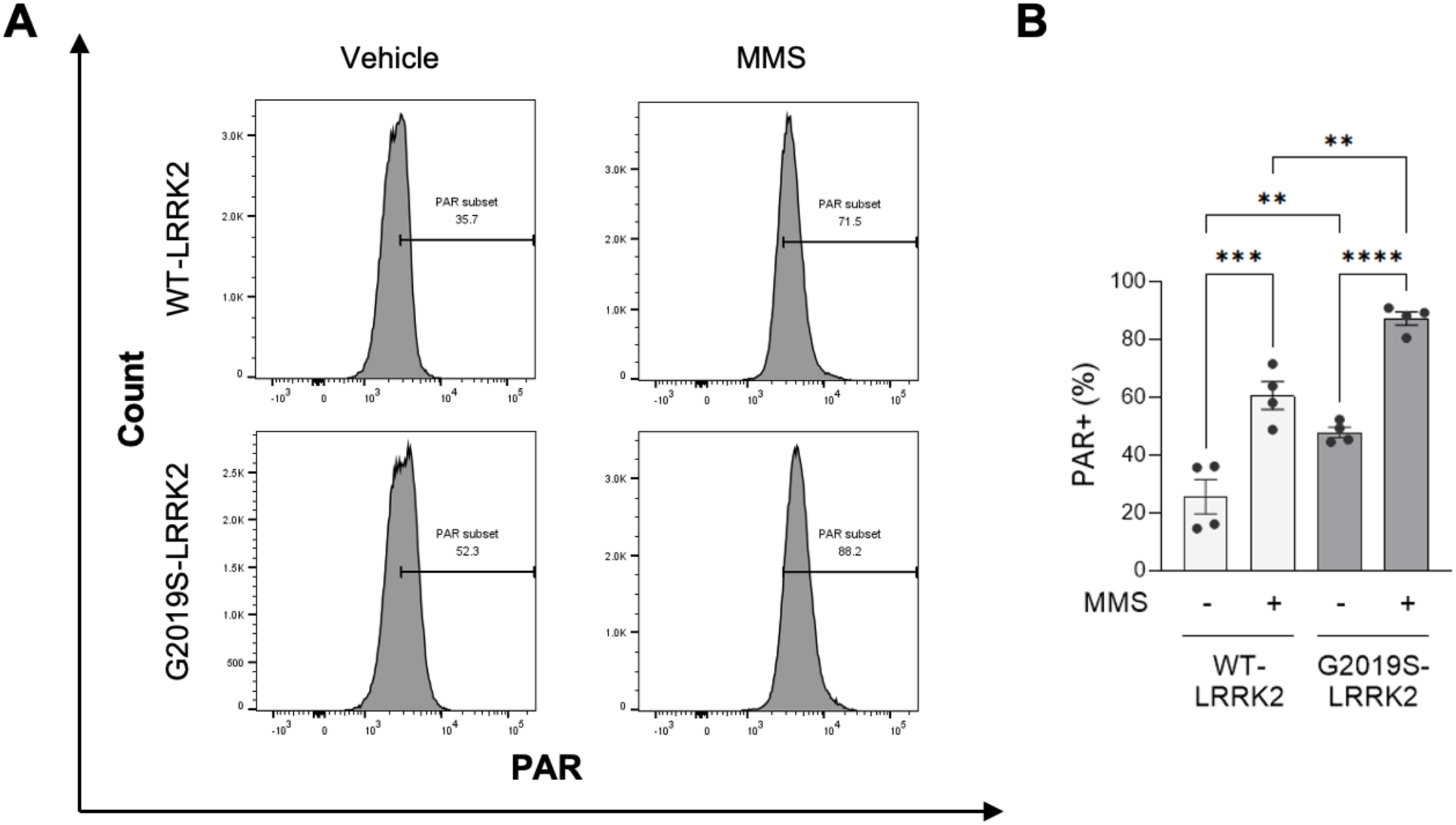
The LRRK2 G2019S mutation sensitizes cells to the alkylating DNA damaging agent *in vitro*. (**A**) Representative flow cytometry plots of WT-LRRK2 and G2019S-LRRK2 expressing cells treated with vehicle or methyl methanesulfonate (MMS, 0.5 µM for 30 min). (**B**) Quantification of PAR-positive cells with and without exposure to MMS. (n=4, **p < 0.01, ***p < 0.001, ****p < 0.0001, determined by one-way ANOVA.) Data are presented as mean ± SEM.

**Supplementary Figure 12.**
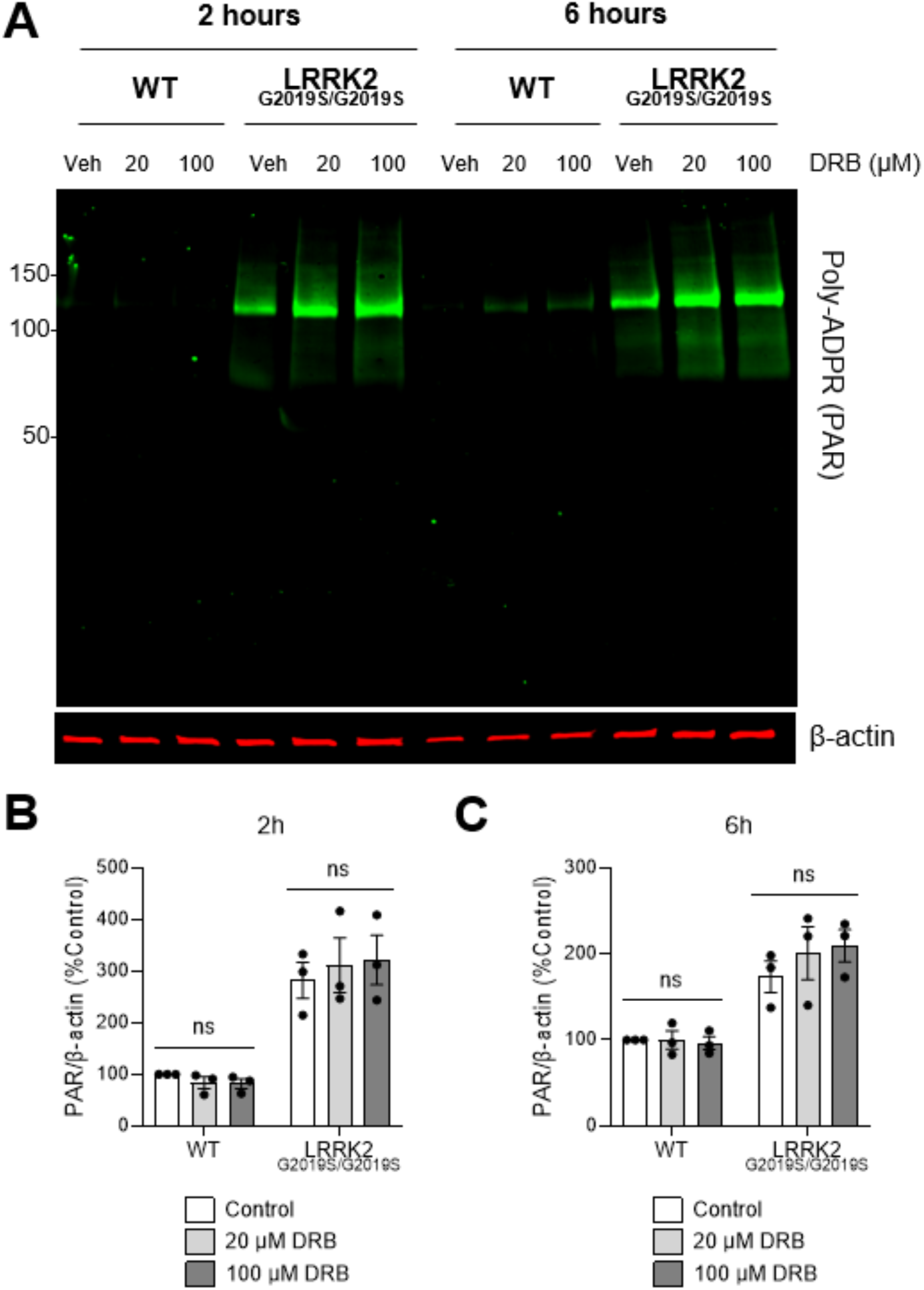
LRRK2 G2019S-mediated PAR accumulation *in vitro* is not driven by transcription-related processes. (**A**) Representative western blots of wild-type (WT) and LRRK2^G2019S/G2019S^ KI cells after treatment with vehicle or RNA Polymerase II inhibition with 5,6-dichloro-1-beta-D-ribofuranosylbenzimidazole (DRB) (20 and 100 µM) for 2 or 6 h and assessed for PAR levels and β-actin as a loading control. (**B**) Quantification of PAR levels were unchanged independent of genotype with either a 2 h treatment or (**C**) 6 h treatment. (n=3, ns, determined by two-way ANOVA with Bonferroni’s multiple comparison). Data are presented as mean ± SEM.

## Notes

### Competing Interest Statement

The authors have declared no competing interest.

## References

1. Lewis, S.J., Gangadharan, S. and Padmakumar, C.P. (2016) Parkinson’s disease in the older patient. Clin Med (Lond), 16, 376–378.

2. Brakedal, B., Toker, L., Haugarvoll, K. and Tzoulis, C. (2022) A nationwide study of the incidence, prevalence and mortality of Parkinson’s disease in the Norwegian population. npj Parkinson’s Disease, 8, 19.

3. Marras, C., Beck, J.C., Bower, J.H., Roberts, E., Ritz, B., Ross, G.W., Abbott, R.D., Savica, R., Van Den Eeden, S.K., Willis, A.W. et al. (2018) Prevalence of Parkinson’s disease across North America. npj Parkinson’s Disease, 4, 21.

4. Yang, W., Hamilton, J.L., Kopil, C., Beck, J.C., Tanner, C.M., Albin, R.L., Ray Dorsey, E., Dahodwala, N., Cintina, I., Hogan, P. et al. (2020) Current and projected future economic burden of Parkinson’s disease in the U.S. npj Parkinson’s Disease, 6, 15.

5. Bloem, B.R., Okun, M.S. and Klein, C. (2021) Parkinson’s disease. The Lancet, 397, 2284–2303.

6. Morris, H.R., Spillantini, M.G., Sue, C.M. and Williams-Gray, C.H. (2024) The pathogenesis of Parkinson’s disease. The Lancet, 403, 293–304.

7. Surmeier, D.J., Obeso, J.A. and Halliday, G.M. (2017) Selective neuronal vulnerability in Parkinson disease. Nat Rev Neurosci, 18, 101–113.

8. Blauwendraat, C., Nalls, M.A. and Singleton, A.B. (2020) The genetic architecture of Parkinson’s disease. Lancet Neurol, 19, 170–178.

9. Thomas, B. and Beal, M.F. (2007) Parkinson’s disease. Hum Mol Genet, 16 Spec No. 2, R183–194.

10. Tran, J., Anastacio, H. and Bardy, C. (2020) Genetic predispositions of Parkinson’s disease revealed in patient-derived brain cells. npj Parkinson’s Disease, 6, 8.

11. Tolosa, E., Vila, M., Klein, C. and Rascol, O. (2020) LRRK2 in Parkinson disease: challenges of clinical trials. Nature Reviews Neurology, 16, 97–107.

12. Healy, D.G., Falchi, M., O’Sullivan, S.S., Bonifati, V., Durr, A., Bressman, S., Brice, A., Aasly, J., Zabetian, C.P., Goldwurm, S. et al. (2008) Phenotype, genotype, and worldwide genetic penetrance of LRRK2-associated Parkinson’s disease: a case-control study. Lancet Neurol, 7, 583–590.

13. Paisán-Ruiz, C., Lewis, P.A. and Singleton, A.B. (2013) LRRK2: Cause, Risk, and Mechanism. Journal of Parkinson’s Disease, 3, 85–103.

14. Tropea, T.F., Hartstone, W., Amari, N., Baum, D., Rick, J., Suh, E., Zhang, H., Paul, R.A., Han, N., Zack, R. et al. (2024) Genetic and phenotypic characterization of Parkinson’s disease at the clinic-wide level. NPJ Parkinsons Dis, 10, 97.

15. Biskup, S. and West, A.B. (2009) Zeroing in on LRRK2-linked pathogenic mechanisms in Parkinson’s disease. Biochim Biophys Acta, 1792, 625–633.

16. Taymans, J.-M., Fell, M., Greenamyre, T., Hirst, W.D., Mamais, A., Padmanabhan, S., Peter, I., Rideout, H. and Thaler, A. (2023) Perspective on the current state of the LRRK2 field. npj Parkinson’s Disease, 9, 104.

17. Sosero, Y.L. and Gan-Or, Z. (2023) LRRK2 and Parkinson’s disease: from genetics to targeted therapy. Ann Clin Transl Neurol, 10, 850–864.

18. Buck, S.A. and Sanders, L.H. (2025) LRRK2-mediated mitochondrial dysfunction in Parkinson’s disease. Biochemical Journal, 482, 721–739.

19. Bender, A., Krishnan, K.J., Morris, C.M., Taylor, G.A., Reeve, A.K., Perry, R.H., Jaros, E., Hersheson, J.S., Betts, J., Klopstock, T. et al. (2006) High levels of mitochondrial DNA deletions in substantia nigra neurons in aging and Parkinson disease. Nat Genet, 38, 515–517.

20. Dölle, C., Flønes, I., Nido, G.S., Miletic, H., Osuagwu, N., Kristoffersen, S., Lilleng, P.K., Larsen, J.P., Tysnes, O.-B., Haugarvoll, K. et al. (2016) Defective mitochondrial DNA homeostasis in the substantia nigra in Parkinson disease. Nature Communications, 7, 13548.

21. Mandavilli, B.S., Ali, S.F. and Van Houten, B. (2000) DNA damage in brain mitochondria caused by aging and MPTP treatment. Brain Res, 885, 45–52.

22. Sanders, L.H., McCoy, J., Hu, X., Mastroberardino, P.G., Dickinson, B.C., Chang, C.J., Chu, C.T., Van Houten, B. and Greenamyre, J.T. (2014) Mitochondrial DNA damage: molecular marker of vulnerable nigral neurons in Parkinson’s disease. Neurobiol Dis, 70, 214–223.

23. Kraytsberg, Y., Kudryavtseva, E., McKee, A.C., Geula, C., Kowall, N.W. and Khrapko, K. (2006) Mitochondrial DNA deletions are abundant and cause functional impairment in aged human substantia nigra neurons. Nat Genet, 38, 518–520.

24. Alam, Z.I., Jenner, A., Daniel, S.E., Lees, A.J., Cairns, N., Marsden, C.D., Jenner, P. and Halliwell, B. (1997) Oxidative DNA damage in the parkinsonian brain: an apparent selective increase in 8-hydroxyguanine levels in substantia nigra. J Neurochem, 69, 1196–1203.

25. Lin, M.T., Cantuti-Castelvetri, I., Zheng, K., Jackson, K.E., Tan, Y.B., Arzberger, T., Lees, A.J., Betensky, R.A., Beal, M.F. and Simon, D.K. (2012) Somatic mitochondrial DNA mutations in early Parkinson and incidental Lewy body disease. Ann Neurol, 71, 850–854.

26. Qi, R., Sammler, E., Gonzalez-Hunt, C.P., Barraza, I., Pena, N., Rouanet, J.P., Naaldijk, Y., Goodson, S., Fuzzati, M., Blandini, F. et al. (2023) A blood-based marker of mitochondrial DNA damage in Parkinson’s disease. Science Translational Medicine, 15, eabo1557.

27. Pena, N., Richbourg, T., Gonzalez-Hunt, C.P., Qi, R., Wren, P., Barlow, C., Shanks, N.F., Carlisle, H.J. and Sanders, L.H. (2024) G2019S selective LRRK2 kinase inhibitor abrogates mitochondrial DNA damage. npj Parkinson’s Disease, 10, 49.

28. Howlett, E.H., Jensen, N., Belmonte, F., Zafar, F., Hu, X., Kluss, J., Schule, B., Kaufman, B.A., Greenamyre, J.T. and Sanders, L.H. (2017) LRRK2 G2019S-induced mitochondrial DNA damage is LRRK2 kinase dependent and inhibition restores mtDNA integrity in Parkinson’s disease. Hum Mol Genet.

29. Sanders, L.H., Laganiere, J., Cooper, O., Mak, S.K., Vu, B.J., Huang, Y.A., Paschon, D.E., Vangipuram, M., Sundararajan, R., Urnov, F.D. et al. (2014) LRRK2 mutations cause mitochondrial DNA damage in iPSC-derived neural cells from Parkinson’s disease patients: reversal by gene correction. Neurobiol Dis, 62, 381–386.

30. Gonzalez-Hunt, C.P., Thacker, E.A., Toste, C.M., Boularand, S., Deprets, S., Dubois, L. and Sanders, L.H. (2020) Mitochondrial DNA damage as a potential biomarker of LRRK2 kinase activity in LRRK2 Parkinson’s disease. Sci Rep, 10, 17293.

31. Guo, J.D., Zhao, X., Li, Y., Li, G.R. and Liu, X.L. (2018) Damage to dopaminergic neurons by oxidative stress in Parkinson’s disease (Review). Int J Mol Med, 41, 1817–1825.

32. Ionescu-Tucker, A. and Cotman, C.W. (2021) Emerging roles of oxidative stress in brain aging and Alzheimer’s disease. Neurobiology of Aging, 107, 86–95.

33. Di Maio, R., Hoffman, E.K., Rocha, E.M., Keeney, M.T., Sanders, L.H., De Miranda, B.R., Zharikov, A., Van Laar, A., Stepan, A.F., Lanz, T.A., et al. (2018) LRRK2 activation in idiopathic Parkinson’s disease. Sci Transl Med, 10.

34. Sanders, L.H. and Timothy Greenamyre, J. (2013) Oxidative damage to macromolecules in human Parkinson disease and the rotenone model. Free Radic Biol Med, 62, 111–120.

35. Gorbunova, V., Seluanov, A., Mao, Z. and Hine, C. (2007) Changes in DNA repair during aging. Nucleic Acids Res, 35, 7466–7474.

36. Chaudhary, M.R., Chaudhary, S., Sharma, Y., Singh, T.A., Mishra, A.K., Sharma, S. and Mehdi, M.M. (2023) Aging, oxidative stress and degenerative diseases: mechanisms, complications and emerging therapeutic strategies. Biogerontology, 24, 609–662.

37. Maynard, S., Fang, E.F., Scheibye-Knudsen, M., Croteau, D.L. and Bohr, V.A. (2015) DNA Damage, DNA Repair, Aging, and Neurodegeneration. Cold Spring Harbor Perspectives in Medicine, 5, a025130–a025130.

38. Hou, Y., Dan, X., Babbar, M., Wei, Y., Hasselbalch, S.G., Croteau, D.L. and Bohr, V.A. (2019) Ageing as a risk factor for neurodegenerative disease. Nat Rev Neurol, 15, 565–581.

39. Lombard, D.B., Chua, K.F., Mostoslavsky, R., Franco, S., Gostissa, M. and Alt, F.W. (2005) DNA Repair, Genome Stability, and Aging. Cell, 120, 497–512.

40. Yousefzadeh, M., Henpita, C., Vyas, R., Soto-Palma, C., Robbins, P. and Niedernhofer, L. (2021) DNA damage—how and why we age? eLife, 10.

41. David, S.S., O’Shea, V.L. and Kundu, S. (2007) Base-excision repair of oxidative DNA damage. Nature, 447, 941–950.

42. Sanders, L.H., Paul, K.C., Howlett, E.H., Lawal, H., Boppana, S., Bronstein, J.M., Ritz, B. and Greenamyre, J.T. (2017) Editor’s Highlight: Base Excision Repair Variants and Pesticide Exposure Increase Parkinson’s Disease Risk. Toxicol Sci, 158, 188–198.

43. Krokan, H.E. and Bjørås, M. (2013) Base excision repair. Cold Spring Harb Perspect Biol, 5, a012583.

44. Hegde, M.L., Hazra, T.K. and Mitra, S. (2008) Early steps in the DNA base excision/single-strand interruption repair pathway in mammalian cells. Cell Research, 18, 27–47.

45. Wu, W., Kargbo-Hill, S.E., Nathan, W.J., Paiano, J., Callen, E., Wang, D., Shinoda, K., van Wietmarschen, N., Colón-Mercado, J.M., Zong, D., et al. (2021) Neuronal enhancers are hotspots for DNA single-strand break repair. Nature, 593, 440–444.

46. Yu, H., Su, Y., Shin, J., Zhong, C., Guo, J.U., Weng, Y.L., Gao, F., Geschwind, D.H., Coppola, G., Ming, G.L. et al. (2015) Tet3 regulates synaptic transmission and homeostatic plasticity via DNA oxidation and repair. Nat Neurosci, 18, 836–843.

47. Madabhushi, R., Pan, L. and Tsai, L.-H. (2014) DNA Damage and Its Links to Neurodegeneration. Neuron, 83, 266–282.

48. Delint-Ramirez, I. and Madabhushi, R. (2025) DNA damage and its links to neuronal aging and degeneration. Neuron, 113, 7–28.

49. Ray Chaudhuri, A. and Nussenzweig, A. (2017) The multifaceted roles of PARP1 in DNA repair and chromatin remodelling. Nature Reviews Molecular Cell Biology, 18, 610–621.

50. Curtin, N.J. and Szabo, C. (2020) Poly(ADP-ribose) polymerase inhibition: past, present and future. Nature Reviews Drug Discovery, 19, 711–736.

51. Laspata, N., Kaur, P., Mersaoui, Sofiane Y., Muoio, D., Liu, Zhiyan S., Bannister, M.H., Nguyen, Hai D., Curry, C., Pascal, John M., Poirier, Guy G. et al. (2023) PARP1 associates with R-loops to promote their resolution and genome stability. Nucleic Acids Research, gkad066.

52. Rouleau, M., Patel, A., Hendzel, M.J., Kaufmann, S.H. and Poirier, G.G. (2010) PARP inhibition: PARP1 and beyond. Nature Reviews Cancer, 10, 293–301.

53. Caldecott, K.W. (2023) Causes and consequences of DNA single-strand breaks. Trends in Biochemical Sciences.

54. Laspata, N., Muoio, D. and Fouquerel, E. (2024) Multifaceted Role of PARP1 in Maintaining Genome Stability Through Its Binding to Alternative DNA Structures. Journal of Molecular Biology, 436, 168207.

55. Salemi, M., Mazzetti, S., De Leonardis, M., Giampietro, F., Medici, V., Poloni, T.E., Cannarella, R., Giaccone, G., Pezzoli, G., Cappelletti, G., et al. (2021) Poly (ADP-ribose) polymerase 1 and Parkinson’s disease: A study in post-mortem human brain. Neurochemistry International, 144, 104978.

56. Maiuri, T., Bazan, C.B., Harding, R.J., Begeja, N., Kam, T.I., Byrne, L.M., Rodrigues, F.B., Warner, M.M., Neuman, K., Mansoor, M. et al. (2024) Poly ADP-ribose signaling is dysregulated in Huntington disease. Proc Natl Acad Sci U S A, 121, e2318098121.

57. Mao, K. and Zhang, G. (2022) The role of PARP1 in neurodegenerative diseases and aging. The FEBS Journal, 289, 2013–2024.

58. Woolley, P.R., Wen, X., Conway, O.M., Ender, N.A., Lee, J.-H. and Paull, T.T. (2024) Regulation of transcription patterns, poly(ADP-ribose), and RNA-DNA hybrids by the ATM protein kinase. Cell Reports, 43, 113896.

59. Kam, T.-I., Mao, X., Park, H., Chou, S.-C., Karuppagounder, S.S., Umanah, G.E., Yun, S.P., Brahmachari, S., Panicker, N., Chen, R. et al. (2018) Poly(ADP-ribose) drives pathologic α-synuclein neurodegeneration in Parkinson’s disease. Science, 362, eaat8407.

60. Jhaldiyal, A., Kumari, M., Tripathi, T., Khan, M.R., Wang, J., Guttman, L., Biswas, D., Pasupuleti, A., Aggarwal, A., Pandya, S. et al. (2025) Poly(ADP-ribose) Polymerase 1 Deficiency Attenuates Amyloid Pathology, Neurodegeneration, and Cognitive Decline in a Familial Alzheimer’s Disease Model. bioRxiv.

61. Yue, M., Hinkle, K.M., Davies, P., Trushina, E., Fiesel, F.C., Christenson, T.A., Schroeder, A.S., Zhang, L., Bowles, E., Behrouz, B. et al. (2015) Progressive dopaminergic alterations and mitochondrial abnormalities in LRRK2 G2019S knock-in mice. Neurobiology of Disease, 78, 172–195.

62. Lozen, M., Chen, Y., Boisvert, R.A. and Schlacher, K. (2023) Mitochondrial Replication Assay (MIRA) for Efficient in situ Quantification of Nascent mtDNA and Protein Interactions with Nascent mtDNA (mitoSIRF). Bio-protocol, 13, e4680.

63. Luzwick, J.W., Dombi, E., Boisvert, R.A., Roy, S., Park, S., Kunnimalaiyaan, S., Goffart, S., Schindler, D. and Schlacher, K. (2021) MRE11-dependent instability in mitochondrial DNA fork protection activates a cGAS immune signaling pathway. Sci Adv, 7, eabf9441.

64. Kelley, M.R., Logsdon, D. and Fishel, M.L. (2014) Targeting DNA repair pathways for cancer treatment: what’s new? Future Oncol, 10, 1215–1237.

65. Timson, J. (1975) Hydroxyurea. Mutation Research/Reviews in Genetic Toxicology, 32, 115–131.

66. Hanzlikova, H., Gittens, W., Krejcikova, K., Zeng, Z. and Caldecott, K.W. (2017) Overlapping roles for PARP1 and PARP2 in the recruitment of endogenous XRCC1 and PNKP into oxidized chromatin. Nucleic Acids Res, 45, 2546–2557.

67. Bhandari, S.K., Wiest, N., Sallmyr, A., Du, R. and Tomkinson, Alan E. (2024) Redundant but essential functions of PARP1 and PARP2 in DNA ligase I-independent DNA replication. Nucleic Acids Research, 52, 10341–10354.

68. Chaitanya, G.V., Alexander, J.S. and Babu, P.P. (2010) PARP-1 cleavage fragments: signatures of cell-death proteases in neurodegeneration. Cell Communication and Signaling, 8, 31.

69. Kanev, P.-B., Varhoshkova, S., Georgieva, I., Lukarska, M., Kirova, D., Danovski, G., Stoynov, S. and Aleksandrov, R. (2024) A unified mechanism for PARP inhibitor-induced PARP1 chromatin retention at DNA damage sites in living cells. Cell Reports, 43.

70. Rong, Y., Doctrow, S.R., Tocco, G. and Baudry, M. (1999) EUK-134, a synthetic superoxide dismutase and catalase mimetic, prevents oxidative stress and attenuates kainate-induced neuropathology. Proc Natl Acad Sci U S A, 96, 9897–9902.

71. Baker, K., Marcus, C.B., Huffman, K., Kruk, H., Malfroy, B. and Doctrow, S.R. (1998) Synthetic Combined Superoxide Dismutase/Catalase Mimetics Are Protective as a Delayed Treatment in a Rat Stroke Model: A Key Role for Reactive Oxygen Species in Ischemic Brain Injury. The Journal of Pharmacology and Experimental Therapeutics, 284, 215–221.

72. Testa, C.M., Sherer, T.B. and Greenamyre, J.T. (2005) Rotenone induces oxidative stress and dopaminergic neuron damage in organotypic substantia nigra cultures. Brain Res Mol Brain Res, 134, 109–118.

73. Van Laar, A.D., Webb, K.R., Keeney, M.T., Van Laar, V.S., Zharikov, A., Burton, E.A., Hastings, T.G., Glajch, K.E., Hirst, W.D., Greenamyre, J.T. et al. (2023) Transient exposure to rotenone causes degeneration and progressive parkinsonian motor deficits, neuroinflammation, and synucleinopathy. npj Parkinson’s Disease, 9, 121.

74. Wang, M.J., Huang, H.Y., Chiu, T.L., Chang, H.F. and Wu, H.R. (2019) Peroxiredoxin 5 Silencing Sensitizes Dopaminergic Neuronal Cells to Rotenone via DNA Damage-Triggered ATM/p53/PUMA Signaling-Mediated Apoptosis. Cells, 9.

75. Sanders, L.H., Howlett, E.H., McCoy, J. and Greenamyre, J.T. (2014) Mitochondrial DNA damage as a peripheral biomarker for mitochondrial toxin exposure in rats. Toxicol Sci, 142, 395–402.

76. Chen, Z., Cao, Z., Zhang, W., Gu, M., Zhou, Z.D., Li, B., Li, J., Tan, E.K. and Zeng, L. (2017) LRRK2 interacts with ATM and regulates Mdm2-p53 cell proliferation axis in response to genotoxic stress. Hum Mol Genet, 26, 4494–4505.

77. Koss, D.J., Todd, O., Menon, H., Anderson, Z., Yang, T., Findlay, L., Graham, B., Palmowski, P., Porter, A., Morrice, N. et al. (2025) A reciprocal relationship between markers of genomic DNA damage and alpha-synuclein pathology in dementia with Lewy bodies. Mol Neurodegener, 20, 34.

78. Rose, E.P., Osterberg, V.R., Gorbunova, V. and Unni, V.K. (2024) Alpha-synuclein modulates the repair of genomic DNA double-strand breaks in a DNA-PK(cs)-regulated manner. Neurobiol Dis, 201, 106675.

79. Schaser, A.J., Osterberg, V.R., Dent, S.E., Stackhouse, T.L., Wakeham, C.M., Boutros, S.W., Weston, L.J., Owen, N., Weissman, T.A., Luna, E. et al. (2019) Alpha-synuclein is a DNA binding protein that modulates DNA repair with implications for Lewy body disorders. Scientific Reports, 9, 10919–10919.

80. Vasquez, V., Mitra, J., Hegde, P.M., Pandey, A., Sengupta, S., Mitra, S., Rao, K.S. and Hegde, M.L. (2017) Chromatin-Bound Oxidized α-Synuclein Causes Strand Breaks in Neuronal Genomes in in vitro Models of Parkinson’s Disease. J Alzheimers Dis, 60, S133–s150.

81. Wang, D., Yu, T., Liu, Y., Yan, J., Guo, Y., Jing, Y., Yang, X., Song, Y. and Tian, Y. (2016) DNA damage preceding dopamine neuron degeneration in A53T human α-synuclein transgenic mice. Biochem Biophys Res Commun, 481, 104–110.

82. Milanese, C., Cerri, S., Ulusoy, A., Gornati, S.V., Plat, A., Gabriels, S., Blandini, F., Di Monte, D.A., Hoeijmakers, J.H. and Mastroberardino, P.G. (2018) Activation of the DNA damage response in vivo in synucleinopathy models of Parkinson’s disease. Cell Death Dis, 9, 818.

83. Outeiro, T.F. and Koss, D.J. (2025) Nuclear Alpha-Synuclein: Mechanisms and Implications for Synucleinopathies. Mov Disord.

84. Du, T., Li, G., Zong, Q., Luo, H., Pan, Y. and Ma, K. (2024) Nuclear alpha-synuclein accelerates cell senescence and neurodegeneration. Immun Ageing, 21, 47.

85. El-Saadi, M.W., Tian, X., Grames, M., Ren, M., Keys, K., Li, H., Knott, E., Yin, H., Huang, S. and Lu, X.-H. (2022) Tracing brain genotoxic stress in Parkinson’s disease with a novel single-cell genetic sensor. Science Advances, 8, eabd1700.

86. Khan, S., Delotterie, D.F., Xiao, J., Thangavel, R., Hori, R., Koprich, J., Alway, S.E., McDonald, M.P. and Khan, M.M. (2025) Crosstalk between DNA damage and cGAS-STING immune pathway drives neuroinflammation and dopaminergic neurodegeneration in Parkinson’s disease. Brain, Behavior, and Immunity, 130, 106065.

87. Hegde, M.L., Gupta, V.B., Anitha, M., Harikrishna, T., Shankar, S.K., Muthane, U., Subba Rao, K. and Jagannatha Rao, K.S. (2006) Studies on genomic DNA topology and stability in brain regions of Parkinson’s disease. Arch Biochem Biophys, 449, 143–156.

88. Chen, X., Xie, C., Tian, W., Sun, L., Zheng, W., Hawes, S., Chang, L., Kung, J., Ding, J., Chen, S. et al. (2020) Parkinson’s disease-related Leucine-rich repeat kinase 2 modulates nuclear morphology and genomic stability in striatal projection neurons during aging. Molecular Neurodegeneration, 15, 12.

89. Naaldijk, Y., Fernández, B., Fasiczka, R., Fdez, E., Leghay, C., Croitoru, I., Kwok, J.B., Boulesnane, Y., Vizeneux, A., Mutez, E. et al. (2024) A potential patient stratification biomarker for Parkinsońs disease based on LRRK2 kinase-mediated centrosomal alterations in peripheral blood-derived cells. npj Parkinson’s Disease, 10, 12.

90. Sepe, S., Milanese, C., Gabriels, S., Derks, K.W., Payan-Gomez, C., van, I.W.F., Rijksen, Y.M., Nigg, A.L., Moreno, S., Cerri, S., et al. (2016) Inefficient DNA Repair Is an Aging-Related Modifier of Parkinson’s Disease. Cell Rep, 15, 1866–1875.

91. Lengyel-Zhand, Z., Puentes, L.N. and Mach, R.H. (2022) PARkinson’s: From cellular mechanisms to potential therapeutics. Pharmacol Ther, 230, 107968.

92. Lee, Y., Kang, H.C., Lee, B.D., Lee, Y.I., Kim, Y.P. and Shin, J.H. (2014) Poly (ADP-ribose) in the pathogenesis of Parkinson’s disease. BMB Rep, 47, 424–432.

93. Hastings, L., Sokratian, A., Apicco, D.J., Stanhope, C.M., Smith, L., Hirst, W.D., West, A.B. and Kelly, K. (2022) Evaluation of ABT-888 in the amelioration of α-synuclein fibril-induced neurodegeneration. Brain Communications, 4, fcac042.

94. Mandir, A.S., Przedborski, S., Jackson-Lewis, V., Wang, Z.Q., Simbulan-Rosenthal, C.M., Smulson, M.E., Hoffman, B.E., Guastella, D.B., Dawson, V.L. and Dawson, T.M. (1999) Poly(ADP-ribose) polymerase activation mediates 1-methyl-4-phenyl-1, 2,3,6-tetrahydropyridine (MPTP)-induced parkinsonism. Proc Natl Acad Sci U S A, 96, 5774–5779.

95. Outeiro, T.F., Grammatopoulos, T.N., Altmann, S., Amore, A., Standaert, D.G., Hyman, B.T. and Kazantsev, A.G. (2007) Pharmacological inhibition of PARP-1 reduces alpha-synuclein- and MPP+-induced cytotoxicity in Parkinson’s disease in vitro models. Biochem Biophys Res Commun, 357, 596–602.

96. Wang, Y., An, R., Umanah, G.K., Park, H., Nambiar, K., Eacker, S.M., Kim, B., Bao, L., Harraz, M.M., Chang, C. et al. (2016) A nuclease that mediates cell death induced by DNA damage and poly(ADP-ribose) polymerase-1. Science, 354.

97. Wang, Y., Kim, N.S., Haince, J.F., Kang, H.C., David, K.K., Andrabi, S.A., Poirier, G.G., Dawson, V.L. and Dawson, T.M. (2011) Poly(ADP-ribose) (PAR) binding to apoptosis-inducing factor is critical for PAR polymerase-1-dependent cell death (parthanatos). Sci Signal, 4, ra20.

98. Shadfar, S., Parakh, S., Jamali, M.S. and Atkin, J.D. (2023) Redox dysregulation as a driver for DNA damage and its relationship to neurodegenerative diseases. Translational Neurodegeneration, 12, 18.

99. Currim, F., Tanwar, R., Brown-Leung, J.M., Paranjape, N., Liu, J., Sanders, L.H., Doorn, J.A. and Cannon, J.R. (2024) Selective dopaminergic neurotoxicity modulated by inherent cell-type specific neurobiology. Neurotoxicology, 103, 266–287.

100. Schöndorf, D.C., Ivanyuk, D., Baden, P., Sanchez-Martinez, A., De Cicco, S., Yu, C., Giunta, I., Schwarz, L.K., Di Napoli, G., Panagiotakopoulou, V., et al. (2018) The NAD+ Precursor Nicotinamide Riboside Rescues Mitochondrial Defects and Neuronal Loss in iPSC and Fly Models of Parkinson’s Disease. Cell Rep, 23, 2976–2988.

101. Brakedal, B., Dölle, C., Riemer, F., Ma, Y., Nido, G.S., Skeie, G.O., Craven, A.R., Schwarzlmüller, T., Brekke, N., Diab, J. et al. (2022) The NADPARK study: A randomized phase I trial of nicotinamide riboside supplementation in Parkinson’s disease. Cell Metab, 34, 396–407.e396.

102. Inzelberg, R., Cohen, O.S., Aharon-Peretz, J., Schlesinger, I., Gershoni-Baruch, R., Djaldetti, R., Nitsan, Z., Ephraty, L., Tunkel, O., Kozlova, E. et al. (2012) The LRRK2 G2019S mutation is associated with Parkinson disease and concomitant non-skin cancers. Neurology, 78, 781–786.

103. Warø, B.J. and Aasly, J.O. (2018) Exploring cancer in LRRK2 mutation carriers and idiopathic Parkinson’s disease. Brain Behav, 8, e00858.

104. Agalliu, I., San Luciano, M., Mirelman, A., Giladi, N., Waro, B., Aasly, J., Inzelberg, R., Hassin-Baer, S., Friedman, E., Ruiz-Martinez, J. et al. (2015) Higher Frequency of Certain Cancers in LRRK2 G2019S Mutation Carriers With Parkinson Disease: A Pooled Analysis. JAMA Neurology, 72, 58–65.

105. Lord, C.J. and Ashworth, A. (2017) PARP inhibitors: Synthetic lethality in the clinic. Science, 355, 1152–1158.

106. Murai, J., Zhang, Y., Morris, J., Ji, J., Takeda, S., Doroshow, J.H. and Pommier, Y. (2014) Rationale for poly(ADP-ribose) polymerase (PARP) inhibitors in combination therapy with camptothecins or temozolomide based on PARP trapping versus catalytic inhibition. J Pharmacol Exp Ther, 349, 408–416.

107. Southgate, H.E.D., Chen, L., Tweddle, D.A. and Curtin, N.J. (2020) ATR Inhibition Potentiates PARP Inhibitor Cytotoxicity in High Risk Neuroblastoma Cell Lines by Multiple Mechanisms. Cancers (Basel), 12.

108. Zatreanu, D., Robinson, H.M.R., Alkhatib, O., Boursier, M., Finch, H., Geo, L., Grande, D., Grinkevich, V., Heald, R.A., Langdon, S. et al. (2021) Polθ inhibitors elicit BRCA-gene synthetic lethality and target PARP inhibitor resistance. Nat Commun, 12, 3636.

109. Bryant, H.E. and Helleday, T. (2006) Inhibition of poly (ADP-ribose) polymerase activates ATM which is required for subsequent homologous recombination repair. Nucleic Acids Res, 34, 1685–1691.

110. Murai, J., Huang, S.Y., Das, B.B., Renaud, A., Zhang, Y., Doroshow, J.H., Ji, J., Takeda, S. and Pommier, Y. (2012) Trapping of PARP1 and PARP2 by Clinical PARP Inhibitors. Cancer Res, 72, 5588–5599.

111. Gottipati, P., Vischioni, B., Schultz, N., Solomons, J., Bryant, H.E., Djureinovic, T., Issaeva, N., Sleeth, K., Sharma, R.A. and Helleday, T. (2010) Poly(ADP-ribose) polymerase is hyperactivated in homologous recombination-defective cells. Cancer Res, 70, 5389–5398.

112. Gassman, N.R., Stefanick, D.F., Kedar, P.S., Horton, J.K. and Wilson, S.H. (2012) Hyperactivation of PARP triggers nonhomologous end-joining in repair-deficient mouse fibroblasts. PLoS One, 7, e49301.

113. Jelezcova, E., Trivedi, R.N., Wang, X.H., Tang, J.B., Brown, A.R., Goellner, E.M., Schamus, S., Fornsaglio, J.L. and Sobol, R.W. (2010) Parp1 activation in mouse embryonic fibroblasts promotes Pol beta-dependent cellular hypersensitivity to alkylation damage. Mutat Res, 686, 57–67.

114. Demin, A.A., Hirota, K., Tsuda, M., Adamowicz, M., Hailstone, R., Brazina, J., Gittens, W., Kalasova, I., Shao, Z., Zha, S. et al. (2021) XRCC1 prevents toxic PARP1 trapping during DNA base excision repair. Molecular Cell, 81, 3018–3030.e3015.

115. Hirota, K., Ooka, M., Shimizu, N., Yamada, K., Tsuda, M., Ibrahim, M.A., Yamada, S., Sasanuma, H., Masutani, M. and Takeda, S. (2022) XRCC1 counteracts poly(ADP ribose)polymerase (PARP) poisons, olaparib and talazoparib, and a clinical alkylating agent, temozolomide, by promoting the removal of trapped PARP1 from broken DNA. Genes Cells, 27, 331–344.

116. Sultana, R., Abdel-Fatah, T., Abbotts, R., Hawkes, C., Albarakati, N., Seedhouse, C., Ball, G., Chan, S., Rakha, E.A., Ellis, I.O. et al. (2013) Targeting XRCC1 deficiency in breast cancer for personalized therapy. Cancer Res, 73, 1621–1634.

117. Ali, R., Alabdullah, M., Alblihy, A., Miligy, I., Mesquita, K.A., Chan, S.Y., Moseley, P., Rakha, E.A. and Madhusudan, S. (2020) PARP1 blockade is synthetically lethal in XRCC1 deficient sporadic epithelial ovarian cancers. Cancer Lett, 469, 124–133.

